# Cortical functional connectivity indexes arousal state during sleep and anesthesia

**DOI:** 10.1101/823963

**Authors:** Matthew I. Banks, Bryan M. Krause, Christopher M. Endemann, Declan I. Campbell, Christopher K. Kovach, M. Eric Dyken, Hiroto Kawasaki, Kirill V. Nourski

**Author notes:** **Corresponding author:** Matthew I. Banks, Ph.D., Professor, Department of Anesthesiology, University of Wisconsin, 1300 University Avenue, Room 4605, Madison, WI 53706.

## Abstract

Disruption of cortical connectivity likely contributes to loss of consciousness (LOC) during both sleep and general anesthesia, but the degree of overlap in the underlying mechanisms is unclear. Both sleep and anesthesia comprise states of varying levels of arousal and consciousness, including states of largely maintained consciousness (sleep: N1, REM; anesthesia: sedated but responsive) as well as states of substantially reduced consciousness (sleep: N2/N3; anesthesia: unresponsive). Here, we tested the hypotheses that (1) cortical connectivity will reflect clear changes when transitioning into states of reduced consciousness, and (2) these changes are similar for arousal states of comparable levels of consciousness during sleep and anesthesia. Using intracranial recordings from five neurosurgical patients, we compared resting state cortical functional connectivity (as measured by weighted phase lag index) in the same subjects across arousal states during natural sleep [wake (WS), N1, N2, N3, REM] and propofol anesthesia [pre-drug wake (WA), sedated/responsive (S) and unresponsive (U)]. In wake states WS and WA, alpha-band connectivity within and between temporal, parietal and occipital regions was dominant. This pattern was largely unchanged in N1, REM and S. Transitions into states of reduced consciousness N2, N3 and U were characterized by dramatic and strikingly similar changes in connectivity, with dominant connections shifting to frontal cortex. We suggest that shifts from temporo-parieto-occipital to frontal cortical connectivity may reflect impaired sensory processing in states of reduced consciousness. The data indicate that functional connectivity can serve as a biomarker of arousal state and suggest common mechanisms of LOC in sleep and anesthesia.

## 1. Introduction

Elucidating the changes in the brain that occur upon loss and recovery of consciousness (LOC, ROC) is critical to our understanding of the neural basis of consciousness, and is a prerequisite for improving diagnosis and prognosis of disorders of consciousness and noninvasive monitoring of awareness in clinical settings (Bayne et al., 2017; Bernat, 2017; Stein and Glick, 2016). A primary hurdle is identifying changes that are specific to LOC and ROC, as opposed to nonspecific changes in brain activity in response to endogenous or exogenous factors (e.g. neuromodulators during sleep or anesthetic agents). This can be clarified by investigating common features of LOC and ROC during sleep and anesthesia (Mashour, 2006; Shushruth, 2013; Tung and Mendelson, 2004). A handful of studies have compared the changes in neural activity that occur during transitions between arousal states during sleep versus anesthesia in human subjects (Li et al., 2018; Murphy et al., 2011), but commonalities in neural mechanisms have been elusive, perhaps because sleep and anesthesia data in these studies were obtained in different subjects, or because of the metrics investigated, or both. Here, we compare changes in functional connectivity in the same subjects during sleep and propofol anesthesia.

Although endogenous sleep and arousal centers play a role in LOC/ROC under both sleep and anesthesia (Lydic and Baghdoyan, 2005), changes in the contents of consciousness are likely secondary to actions in neocortex (Voss et al., 2019), which is the focus of the current study. Common mechanisms for LOC/ROC under sleep and anesthesia are suggested by similar effects of LOC on sensory cortex observed under both conditions. For example, primary sensory cortex is still responsive to environmental stimuli, and basic organizational features such as frequency tuning in auditory cortex are preserved (Nir et al., 2015; Raz et al., 2014), while responses in higher order cortical sensory areas are largely suppressed (Liu et al., 2012; Wilf et al., 2016). In addition, cortical connectivity, which is central to leading theories of consciousness (Dehaene and Changeux, 2011; Friston, 2005; Tononi et al., 2016), is altered upon LOC during anesthesia (Boly et al., 2012a; Lee et al., 2017; Lee et al., 2013b; Murphy et al., 2011; Ranft et al., 2016; Sanders et al., 2018) and non-rapid eye movement (NREM) sleep (Boly et al., 2012b; Spoormaker et al., 2010).

These studies suggest that LOC under a variety of conditions converges on specific changes in cortical connectivity. However, a major impediment to identifying these changes is a lack of consensus on key details, for example whether overall or long-range connectivity decreases (Boly et al., 2012a; Lee et al., 2013b; Ranft et al., 2016; Spoormaker et al., 2010) or increases (Boly et al., 2012b; Lee et al., 2017; Monti et al., 2013; Murphy et al., 2011) upon LOC. Moreover, despite the evidence for common mechanisms of LOC under anesthesia and during NREM sleep, there are obvious differences between sleep and anesthesia as well (Akeju and Brown, 2017). Specifically, subjects are arousable from the latter but not from the former, and this maintained connectedness with the environment likely involves cortical activation. The structure of natural sleep, in its transitions between REM and multiple stages of NREM sleep, is not mimicked by steady-state anesthesia. A recent imaging study found substantial differences in the changes in functional magnetic resonance imaging (fMRI) functional connectivity that occur during sleep and propofol anesthesia (Li et al., 2018). Furthermore, delta-band activity during the deepest stages of NREM sleep (N3) most closely resembles brain activity under anesthesia (Murphy et al., 2011), but unresponsiveness (and presumably reduced level of consciousness) occurs as well in stage 2 NREM (N2) sleep (Strauss et al., 2015). Direct comparisons of changes in connectivity associated with LOC under natural sleep and anesthesia may help resolve these discrepancies.

Here, we investigated changes in cortical functional connectivity across arousal states under natural sleep and anesthesia. Intracranial recordings obtained from neurosurgical patients with pharmacologically resistant epilepsy allowed us to compare connectivity using data obtained from the same recording sites in the same subjects.

## 2. Materials and Methods

### 2.1 Subjects

Experiments were carried out in five neurosurgical patients diagnosed with medically refractory epilepsy who were undergoing chronic invasive electrophysiological monitoring to identify seizure foci prior to resection surgery (Supplementary Table 1). Research protocols were approved by the University of Iowa Institutional Review Board and the National Institutes of Health, and written informed consent was obtained from all subjects. Research participation did not interfere with acquisition of clinically necessary data, and subjects could rescind consent for research without interrupting their clinical management. Subjects were right handed, left language-dominant native English speakers. All subjects underwent standard neuropsychological assessment prior to electrode implantation, and none had cognitive deficits that would impact the results of this study. The subjects were tapered off their antiepileptic medication during chronic monitoring when overnight sleep data were collected (see below). All subjects had their medication regimens reinstated at the end of the monitoring period, prior to induction of general anesthesia for the resection surgery.

### 2.2 Experimental procedures

Electrocorticographic (ECoG) recordings were made using subdural and depth electrodes (Ad-Tech Medical, Racine, WI). Subdural arrays consisted of platinum-iridium discs (2.3 mm diameter, 5-10 mm inter-electrode distance), embedded in a silicon membrane. Depth arrays (8-12 electrodes, 5 mm inter-electrode distance) were stereotactically implanted along the anterolateral-to-posteromedial axis of Heschl’s gyrus (HG). Additional arrays targeted insular cortex and provided coverage of planum temporale and planum polare. This allowed for bracketing suspected epileptogenic zones from dorsal, ventral, medial and lateral aspects (Nagahama et al., 2018; Reddy et al., 2010; Supplementary Fig. 1). Depth electrodes also targeted amygdala and hippocampus, and provided additional coverage of the superior temporal sulcus. A subgaleal electrode was used as a reference. All electrodes were placed solely on the basis of clinical requirements, as determined by the team of epileptologists and neurosurgeons (Nourski and Howard, 2015).

Two sets of no-task, resting-state (RS) data were recorded: overnight sleep data and anesthesia data. RS ECoG, EEG and video data were collected from subjects during natural overnight sleep (Supplementary Fig. 2a). Sleep data were collected in the dedicated, electrically shielded suite in The University of Iowa Clinical Research Unit while the subjects lay in the hospital bed. Data were recorded using a Neuralynx Atlas System (Neuralynx Inc., Bozeman, MT), amplified, filtered (0.1–4000 Hz bandpass, 12 dB/octave rolloff), sampled at 16 kHz. Stages of sleep were defined manually using facial EMG and scalp EEG data based on standard clinical criteria (2017) by board-certified physicians who participate in the inter-scorer reliability program of the AASM. Scalp and facial electrodes were placed by an accredited technician, and data were recorded by a clinical acquisition system (Nihon Kohden EEG-2100) in parallel with research acquisition. Facial electrodes were placed following guidelines of the AASM ^90^ at the left and right mentalis for EMG and adjacent to left and right outer canthi for EOG. EEG was obtained from electrodes placed following the international 10-20 system at A1, A2, F3, F3, F4, O1 and O2 in all subjects, with the following additional electrodes: C3 and C4 in all subjects but R376; E1 and E2 in L372 and R376; CZ and FZ in L409 and L423; and F8 in L423. All subjects had periods of REM, N1 and N2 sleep identified; three out of five subjects had N3 sleep periods as well. One subject (L403) experienced multiple seizures in the second half of the night; those data were excluded from analysis.

Anesthesia RS data were collected in the operating room prior to electrode removal and seizure focus resection surgery. Data were recorded using a TDT RZ2 processor (Tucker-Davis Technologies, Alachua, FL), amplified, filtered (0.7–800 Hz bandpass, 12 dB/octave rolloff), and digitized at a sampling rate of 2034.5 Hz. We note that the highpass cutoff frequency on this hardware precluded analysis of frequencies below 1 Hz. Although no specific instructions were given about keeping eyes open or closed, subjects were observed to have eyes closed during nearly all resting state recordings. Data were recorded in 6-minute blocks, interleaved with an auditory stimulus paradigm as part of a separate study (Nourski et al., 2018a, b). Data were collected during an awake baseline period and during induction of general anesthesia with incrementally titrated propofol infusion (50 – 150 μg/kg/min; Supplementary Fig. 2b).

Awareness was assessed using the Observer’s Assessment of Alertness/Sedation (OAA/S) scale (Chernik et al., 1990), and using the bispectral index [BIS (Gan et al., 1997)] (BIS Complete 4-Channel Monitor; Medtronic) recorded continuously throughout the experiment. OAA/S was assessed just before and just after collection of each RS data block. Two levels of anesthesia (arousal states) were targeted: sedated but responsive to command (S; OAA/S ≥ 3) and unresponsive (U; OAA/S ≤ 2) (Nourski et al., 2018a). In four of five subjects, OAA/S values crossed the boundary between S and U over the course of the 6-minute RS block (e.g. RS block #1 in subject L372; see Supplementary Fig. 2b). In these cases, only the first and last 60-second segments of the block were analyzed; data from the first segment were assigned to the S state, and data from the second segment were assigned to the U state.

### 2.3 Data analysis

#### 2.3.1 Band power analysis

Data were assigned to specific arousal states based on sleep scoring and OAA/S assessment. For each subject, sleep and anesthesia data were divided into segments of length 60 seconds for all analyses except the classification analysis (Fig. 5; see below), for which 10-second segments were used. Time-frequency analysis was performed using the demodulated band transform (DBT; Kovach and Gander, 2016), which optimizes frequency resolution for each frequency band specified, while minimizing spectral leakage across bands. PSDs were estimated for each data segment from the squared magnitude of the DBT. For each subject, PSDs were averaged across segments assigned to identical arousal states. ECoG band power was calculated as the average power across frequency in each band. Band power within ROI group was computed as the average across all recording sites in that ROI group, and arousal state-dependent changes in band power were evaluated using linear mixed effects models as follows. The data were normalized to total power and log transformed, then fit with a model incorporating fixed effects of state, ROI, and the interaction of state and ROI, and random effects for channels nested within subjects and with random slopes for brain state by subject, using the R package lme4 (Bates et al., 2015). Estimated marginal means and 95% CIs for each ROI and state were calculated, as well as pairwise between-states contrasts within each ROI with *p*-values adjusted by multivariate *t* for all comparisons within a band, using the R package emmeans (Lenth, 2019).

#### 2.3.2 Connectivity analysis

Connectivity was measured using the debiased weighted phase lag index (wPLI) (Vinck et al., 2011), a non-directed measure of phase synchronization that eschews synchronization near zero phase lag to avoid artifacts due to volume conduction. For each data segment, wPLI was estimated for every electrode pair from the sign of the imaginary part of the DBT-derived cross-spectrum at each frequency and averaged across frequencies within each band of interest (delta: 1-4 Hz, theta: 4-8 Hz, alpha: 8-13 Hz, beta: 13-30 Hz, gamma: 30-70 Hz; high gamma: 70-120 Hz). As the analysis results tended to be correlated in the frequency domain, we chose to present only the results for the delta, alpha and gamma band. Alpha-band wPLI in particular is a commonly used measure of functional connectivity (Blain-Moraes et al., 2014; Blain-Moraes et al., 2015; Lee et al., 2013a; Lee et al., 2017; van Dellen et al., 2014). In addition, we observed evidence for alpha-band oscillatory components in the resting state power spectra, further motivating focus on this band. Therefore, our primary measure of functional connectivity was alpha-band wPLI, but connectivity in delta and gamma bands is presented as well for comparison.

#### 2.3.3 Anatomical reconstruction and ROI parcellation

Electrode localization relied on post-implantation T1-weighted structural MR images and post-implantation CT images. All images were initially aligned with pre-operative T1 images using linear coregistration implemented in FSL (FLIRT) (Jenkinson et al., 2002). Electrodes were identified in the post-implantation MRI as magnetic susceptibility artifacts and in the CT as metallic hyperdensities. Electrode locations were further refined within the space of the pre-operative MRI using three-dimensional non-linear thin-plate spline warping (Rohr et al., 2001), which corrected for post-operative brain shift and distortion. The warping was constrained with 50-100 control points, manually selected throughout the brain, which aligned to visibly corresponding landmarks in the pre- and post-implantation MRIs.

To compare functional connectivity between arousal states, the dimensionality of the adjacency matrices (i.e. the wPLI connectivity matrices) was reduced by assigning electrodes to one of 37 specific ROIs organized into 7 ROI groups (Fig. 3; Table 1; Supplementary Table 2) based upon anatomical reconstructions of electrode locations in each subject. For subdural arrays, it was informed by automated parcellation of cortical gyri (Destrieux et al., 2010; Destrieux et al., 2017) as implemented in the FreeSurfer software package. For depth arrays, ROI assignment was informed by MRI sections along sagittal, coronal and axial planes. For recording sites in HG, delineation of core auditory cortex and adjacent non-core areas (HGPM and HGAL, respectively) was based on physiological criteria (Brugge et al., 2009; Nourski et al., 2016). Specifically, recording sites were assigned to the HGPM ROI if they exhibited phase-locked ECoG responses to 100 Hz click trains and if the averaged evoked potentials to these stimuli featured short-latency (<20 ms) components. Such response features are not present within HGAL. Additionally, correlation coefficients between average evoked potential waveforms recorded from adjacent sites were examined to identify discontinuities in response profiles along HG that could be interpreted as reflecting a transition from HGPM to HGAL. Recording sites identified as seizure foci or characterized by excessive noise, and depth electrode contacts localized to the white matter or outside brain, were excluded from analyses and are not listed in Supplementary Table 2.

**Fig. 1:**
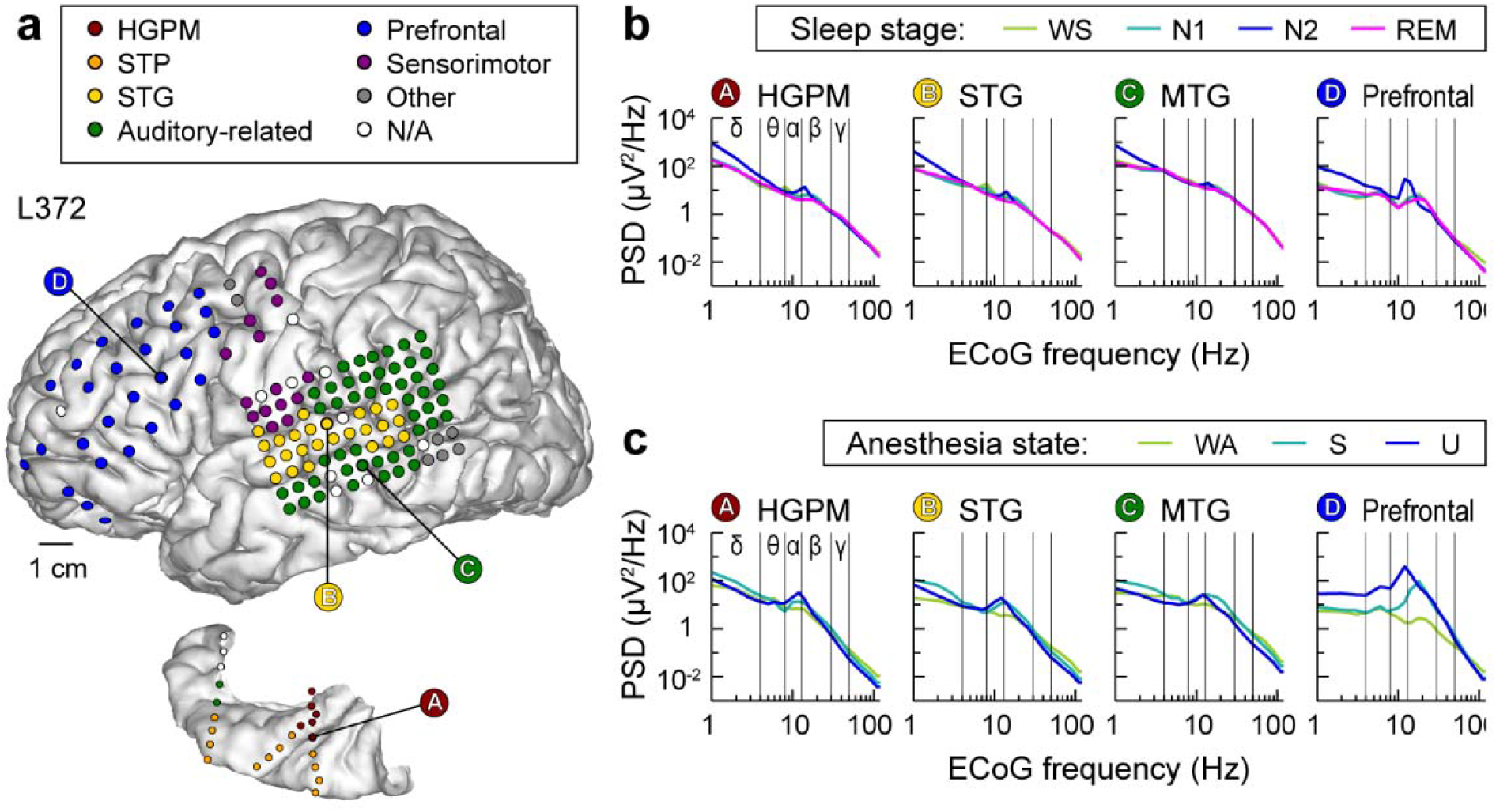
Electrode coverage and electrocorticographic (ECoG) power spectra. Exemplary data from subject L372. **a,** Electrode coverage of the lateral surface of the left cerebral hemisphere (top) and left superior temporal plane (bottom). Recording sites are color-coded according to the region of interest group (see Methods for details and Supplementary table 2 for abbreviation key). **b,** ECoG power spectra during sleep. Data from four representative sites (left-to-right). WS: wake (sleep experiment); PSD: power spectral density. **c,** ECoG power spectra during anesthesia. WA: wake (anesthesia experiment); S: sedated; U: unresponsive.

**Fig. 2:**
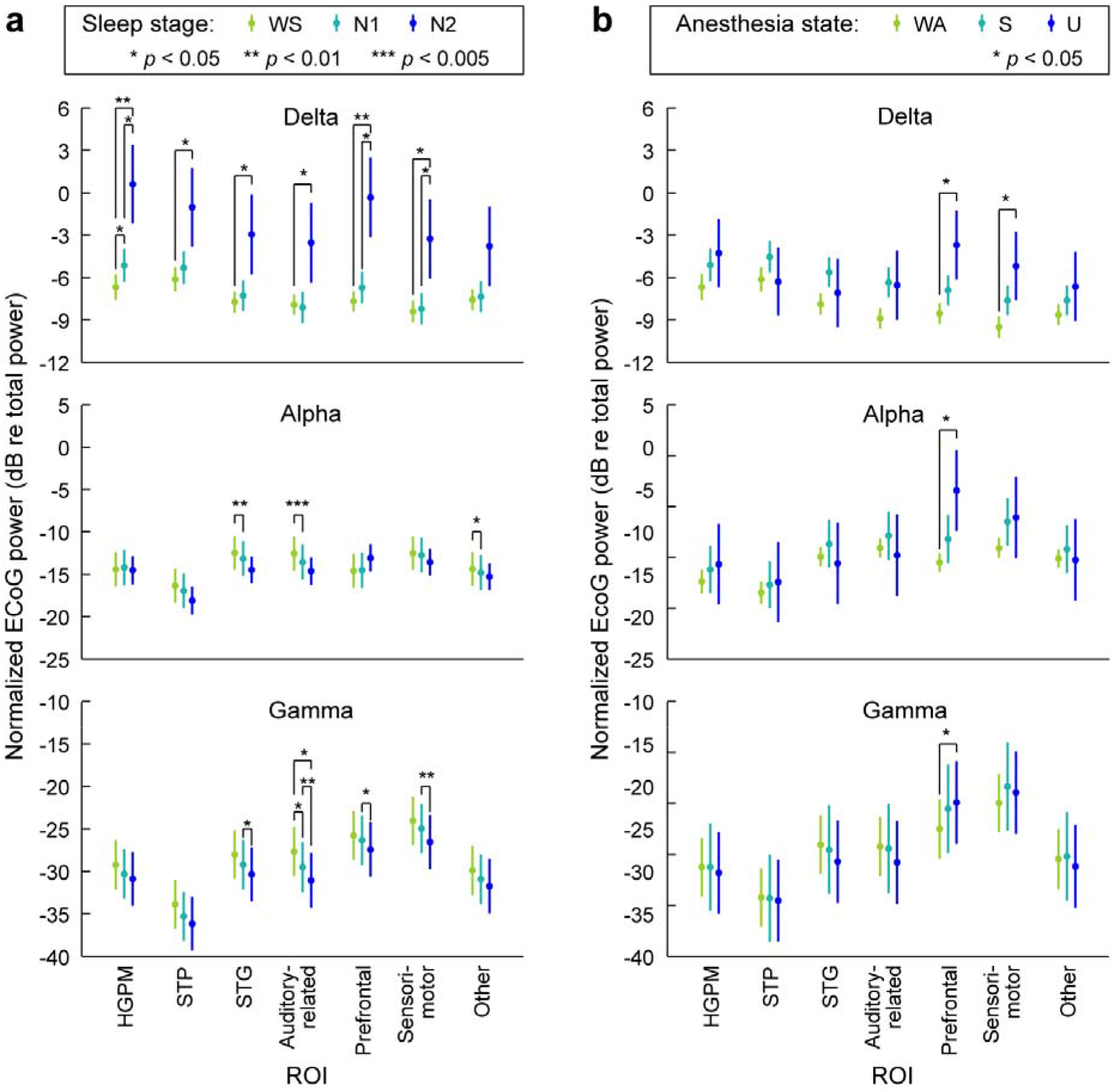
Changes in ECoG band power across arousal states. **a,** ECoG band power during sleep, plotted as marginal means and 95% confidence intervals. **b,** ECoG band power during anesthesia. Data from 5 subjects. Changes in delta, alpha and gamma power are shown in top, middle and bottom rows, respectively. WS: wake (sleep experiment), WA: wake (anesthesia experiment); S: sedated; U: unresponsive.

**Fig. 3:**
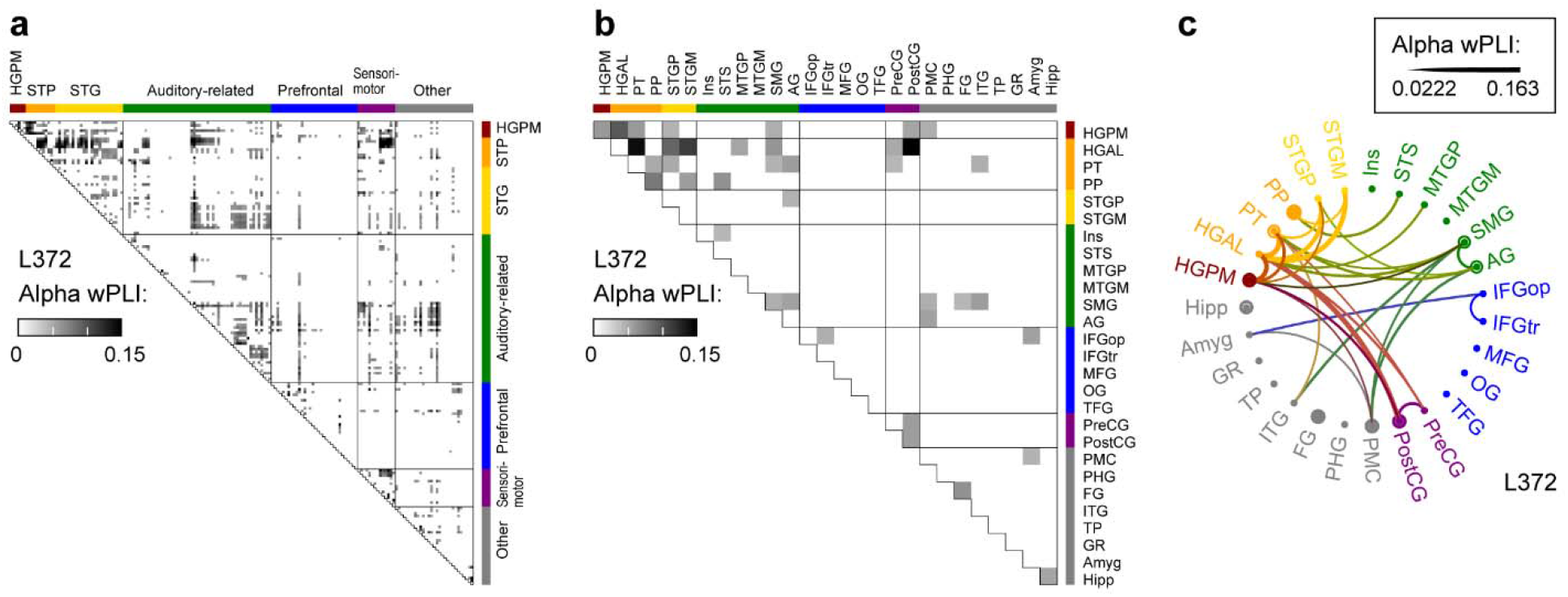
Analysis of alpha-band functional connectivity in wake state. Example from subject L372. **a,** Adjacency matrix for all recording sites. **b,** Adjacency matrix, collapsed for all regions of interest (ROIs). **c,** Chord connectivity plot. Line thickness reflects mean wPLI values that characterize pairs of ROIs. For display purposes, the chord plot was thresholded to retain the 10% strongest connections.

**Table 1.**
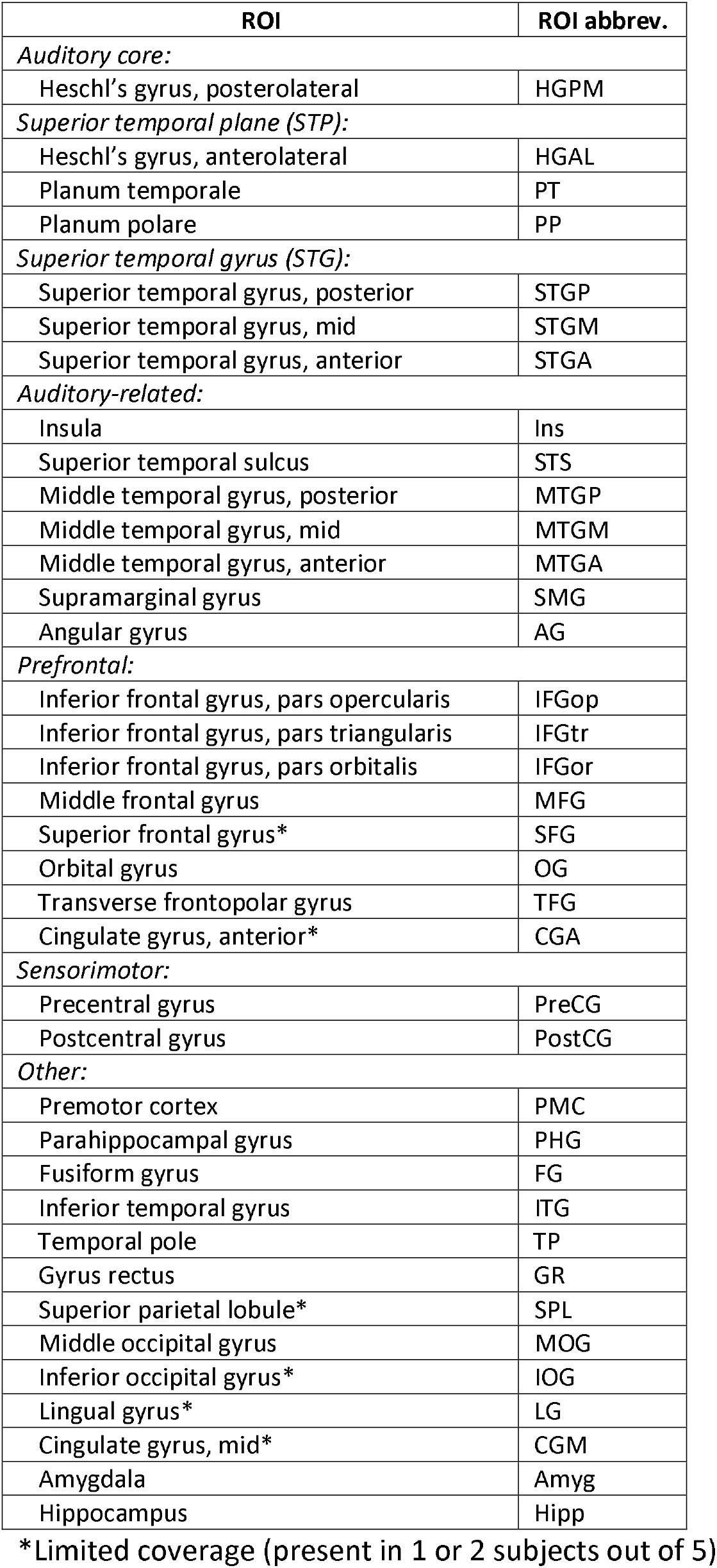
Regions of interest.

#### 2.3.4 ROI-based connectivity analysis

Connectivity between ROIs was computed as the average wPLI value between all pairs of recording sites in the two ROIs. For analyses in which connectivity was summarized across subjects (see Fig. 4 and Supplementary Figs. 6 & 7), ROIs were only included if at least 3 out of 5 subjects had electrode coverage in that ROI; 29 out of 37 ROIs met this criterion. For display purposes only, adjacency matrices for each subject were averaged across segments assigned to identical arousal states, and the matrices thresholded to retain only the 10% strongest connections. Quantitative analyses were based on unthresholded adjacency matrices.

**Fig. 4:**
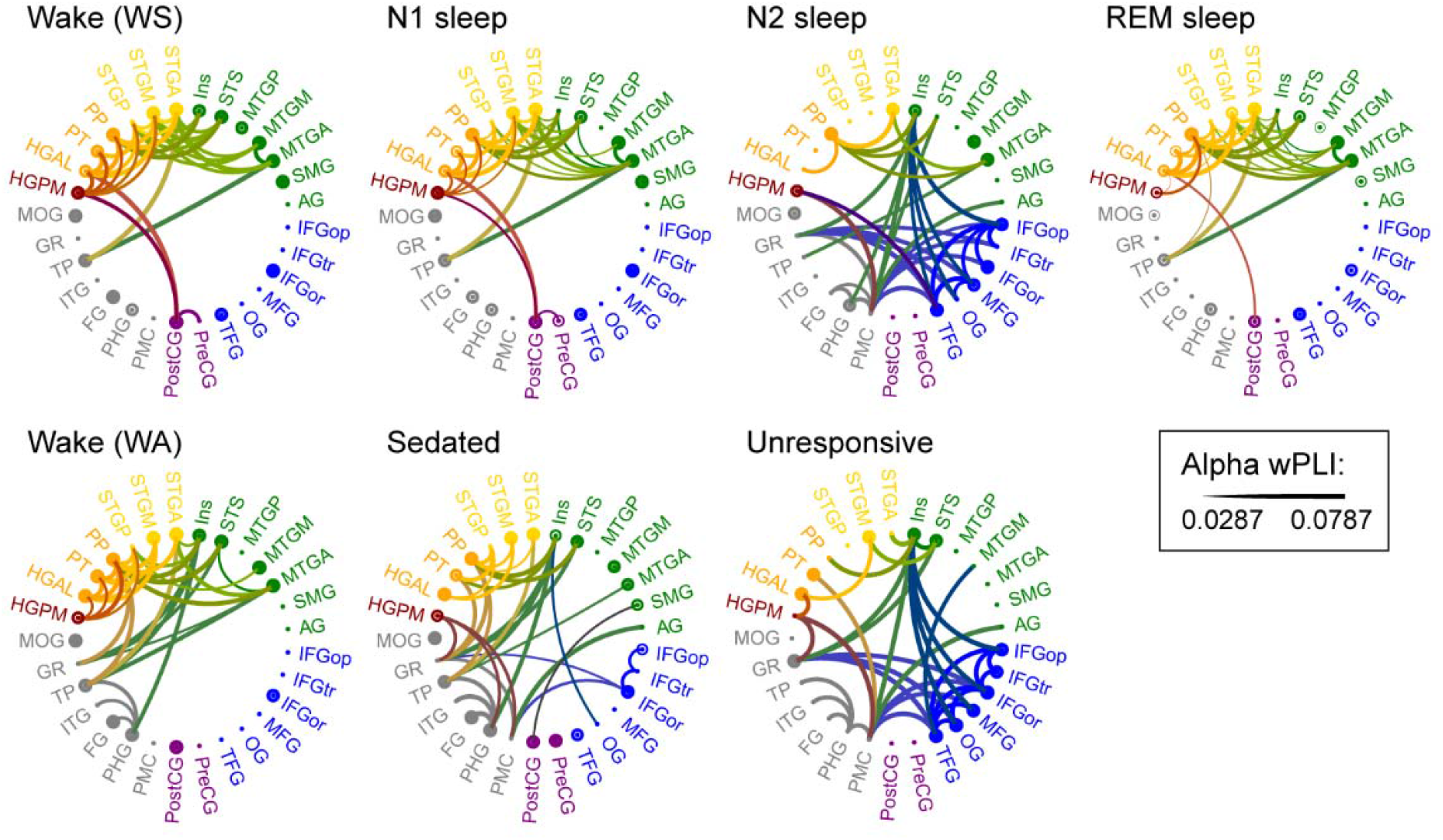
ROI-based analysis of alpha-band functional connectivity across arousal states. Data from five subjects. See caption of **Fig. 3c** for details.

Changes in connectivity with arousal state were evaluated by computing differences between adjacency matrices, and quantified by calculating the operator norm (*d*) of the difference matrix; smaller values of *d* indicate more similar matrices. This difference metric was chosen instead of either the Pearson correlation or the Frobenius norm because it retains information about the structure of the matrix. Specifically, for a matrix **M**, *d*_M_ is the maximum, over all vectors **v** with ∥**v**∥ = 1, of ∥**Mv**∥, and indicates how much **M** stretches these vectors; with **M** representing the difference between adjacency matrices measured in two arousal states, **v** could represent the inputs to or the activity of the nodes of the network at a particular time point, and **Mv** would then be the effect on that activity of the difference in brain state. The operator norm [computed in Matlab as norm (**M**)] is related to the spectrum of **M**^T^**M**: *d*_M_ = the square root of the maximum eigenvalue of **M**^T^**M**.

To compare arousal state-dependent differences in *d* (for example, to see whether d_WS,N1_ is different than *d*_N1,N2_), effect sizes were calculated as Cliff’s delta, δ; (Cliff, 1993). Cliff’s delta ranges from −1 to 1 where 0 indicates completely overlapping distributions and −1 or 1 indicate distributions where all observed values of one group are less/greater than all observed values of the comparison group. Effect sizes were first calculated for each subject, and then reported as the mean effect size across subjects, δ◻. A permutation method was used to estimate *p*-values for these comparisons; within each subject and each experiment (sleep and anesthesia), restricted random permutations of state labels for the data segments, preserving the order of observations, produced an estimated distribution under the null hypothesis that the comparisons do not depend on arousal state (Besag and Clifford, 1989; Winkler et al., 2015). Independent *p*-values obtained within individual subjects for a given test were combined across subjects using Stouffer’s *Z*-transform method (Heard and Rubin-Delanchy, 2018; Stouffer et al., 1949). Non-parametric approaches (Cliff’s delta and permutation method) were preferable to parametric statistics for these data, as the distributions of operator norms and differences were skewed and the magnitude varied between subjects. Given the small number of subjects, these statistical methods treat each subject as a single-case and then combine results in a meta-analysis. Because *p*-values and effect sizes were first estimated in single subjects, this approach reduces the influence of possible outlier subjects and non-normally distributed measures.

#### 2.3.5 Classification analysis

We used a classification analysis as an additional evaluation of changes in connectivity as a function of arousal state. Here, data from each subject was divided into 10-second segments, and adjacency matrices were computed for each segment. To ensure that the data from the two experiments (sleep and anesthesia) were on the same scale, adjacency matrices computed from the anesthesia data were scaled by the slope derived from a regression analysis that related wPLI values computed for sleep vs. anesthesia data for each subject. A linear classifier (implemented using SGDClassifier from Python’s Scikit-Learn library) was trained on a subset (80%) of WS and N2 segments, and then applied to unseen data from all arousal states (WS, N1, N2, N3, REM, WA, S, U) in each subject. Data from the sleep experiment were chosen over those from the anesthesia experiment to train the classifier because the former yielded many more data segments (see Supplementary Fig. 2). Rather than using a binary classification, we applied a logistic weighting function that assigned each segment a weight from 0 (most ‘N2-like’) to 1 (most ‘WS-like’). We report the median logistic prediction scores across all 25 pairwise permutations of WS and N2 train/test splits (4/5 train, 1/5 test) in each subject. Given an unequal number of observations in WS and N2 datasets (see Supplementary Table 3), training sets were balanced in each permutation via random sampling. Hyperparameters corresponding to the strength of regularization (alpha parameter) and the tolerance threshold (i.e. when to stop training the model) were optimized for each training set permutation using three-fold cross-validation. Specifically, each training set was split into three folds, and one of those three folds was used as a test set to evaluate the performance of a given hyperparameter value when training a model on the remaining two folds. For each hyperparameter value evaluated, this process was repeated three times to average over all test sets. Hyperparameter values yielding the lowest average test set error were then used in the final model being applied to unseen data for each train/test permutation. Probability density functions for each arousal state and each subject were estimated from logistic prediction scores using kernel density estimation (ksdensity function in Matlab) and represented as violin plots (see Fig. 5 and Supplementary Fig. 8).

**Fig. 5:**
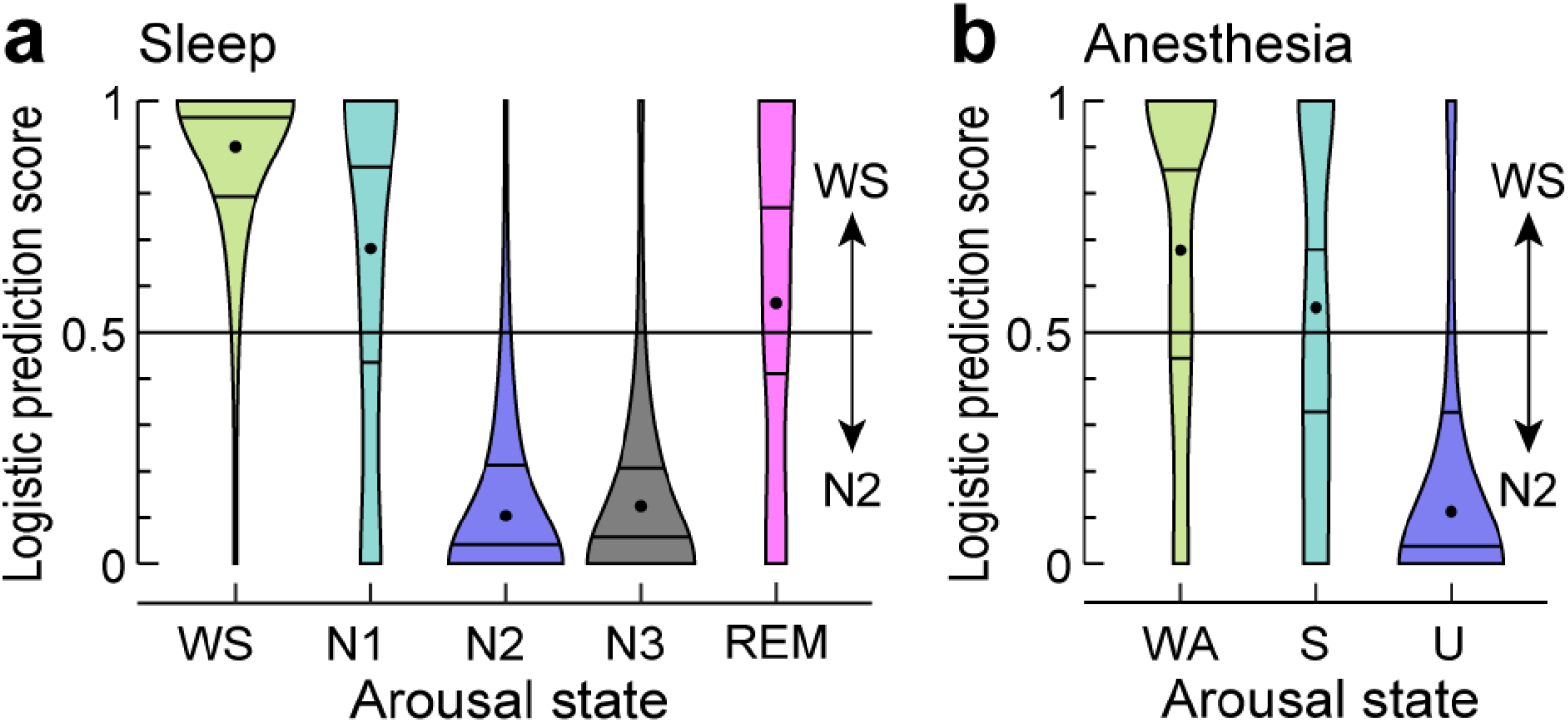
Classification of data segments. Logistic prediction distributions for adjacency matrices from sleep and anesthesia arousal states (panels **a** and **b**, respectively) analyzed by a linear classifier trained on a subset of WS and N2 data. Each violin plot shows the average distribution across five subjects (except for N3, which is for 3 subjects). Centered dot and surrounding horizontal lines represent each distribution’s median and first and third quartiles, respectively. For distributions from individual subjects, see Supplementary Figure 8.

**Fig. 6:**
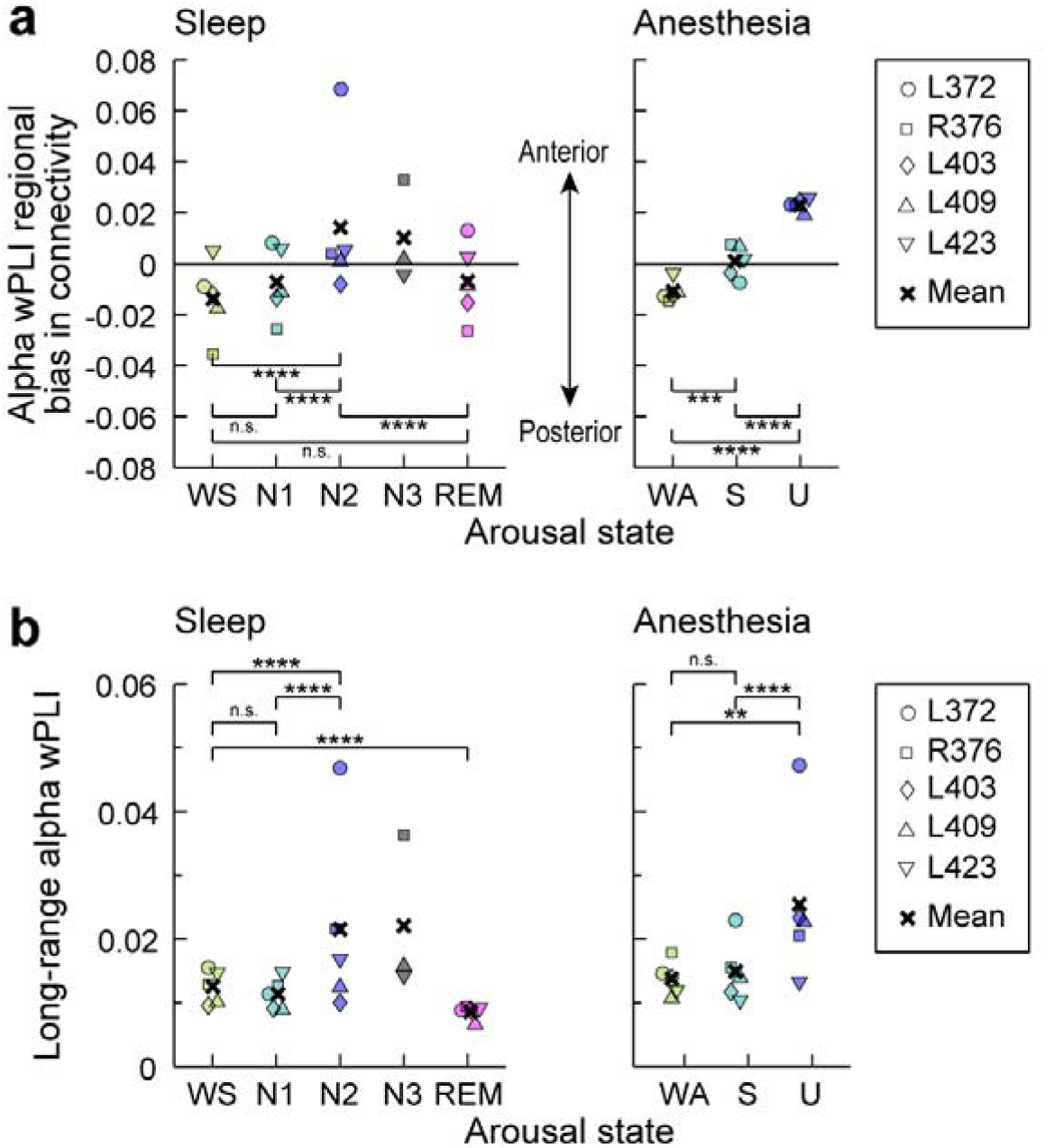
Intra-regional and long-range connectivity changes with arousal state. **a:** Mean alpha wPLI averaged within posterior quadrants of the adjacency matrices minus the average within anterior quadrants. Values greater than zero indicate greater within-posterior connectivity compared to within-anterior connectivity. **b:** Mean alpha wPLI values for recording site pairs distanced greater than the 75th percentile. Significance: n.s., *p* > 0.05; **, *p* < 0.01; ***, *p* < 0.005; ****, *p* < 0.001 (permutation test). Although subject L372 exhibited larger effects than the others in N2 for both analyses, and in U for the long-range connectivity analysis, statistical significance and conclusions were robust to omitting that subject’s (or any individual subject’s) data from the analyses.

#### 2.3.6 Regional connectivity analysis

State-dependent differences in regional connectivity were quantified by dividing ROIs into a posterior (‘back’) group (temporal, parietal and occipital ROIs), and an anterior (‘front’) group (frontal ROIs). Mean alpha-band wPLI across all pairs of recording sites within each group were used to calculate bias in connectivity, defined as the difference between within-posterior and within-anterior connectivity. State-dependence of long-range alpha-band connectivity was assayed by measuring wPLI across the top 25% most distant pairs of recording sites. Euclidean distances between sites were measured using standard 3D coordinates (Right-Anterior-Superior, RAS). Changes in within-posterior versus within-anterior connectivity and changes in long-range connectivity were assessed using permutation analysis as described above for state-dependent differences in *d*.

## 3. Results

### 3.1 Electrode coverage

Data from a total of 864 recording sites from five subjects (Supplementary Table 1), spanning a total of 37 regions of interest (ROIs) were analyzed (Table 1). Each subject contributed between 154 and 198 sites (median 172; Supplementary Table 2, Supplementary Fig. 1). The focus of this study was on changes in cortical connectivity across arousal states. As sensory awareness is a key element of consciousness (Boly et al., 2017), we centered our analysis around cortical hierarchical organization in the auditory modality, which is a convenient choice and a frequent focus of studies of both sleep and general anesthesia (e.g. Liu et al., 2012; Raz et al., 2014; Strauss et al., 2015). Clinical considerations dictated dense sampling of the temporal lobe, including auditory and auditory-related cortex, providing comprehensive electrode coverage across multiple levels of the auditory cortical hierarchy in all subjects.

### 3.2 Defining arousal states

Polysomnography based on scalp electroencephalography (EEG), electrooculography, electromyography, and video was used to assign sleep stages. All five subjects exhibited overnight sleep patterns typical of healthy adult subjects (Supplementary Fig. 2a). There was a high correspondence between the ratio of delta to beta band power in frontal ECoG electrodes and the assigned sleep stage (cf. Kremen et al., 2019). Overnight recordings in all subjects featured wake (WS) state as well as N1, N2 and REM sleep stages; N3 was also observed in 3 of 5 subjects (Supplementary Table 3). The total duration of scored recordings in each subject was between 306.8 and 649.6 minutes (median 534.4).

During the anesthesia experiment, all subjects transitioned from wake (WA) to sedated (S; OAA/S>2) and unresponsive (U; OAA/S ≤2) states as propofol infusion rate was increased (Supplementary Fig. 2b). OAA/S scores exhibited a good correspondence with bispectral index (BIS) values, as expected for sedation and anesthesia induced by propofol alone (Glass et al., 1997). WA, S and U states were characterized by median BIS values of 93 (range 80-98), 78 (range 36-97) and 52 (range 33-74), respectively.

### 3.3 Changes in spectral power under sleep and anesthesia

Power spectral density (PSD) measurements made during WS and WA states exhibited shapes typical of resting state eyes-closed recordings, with power falling off approximately as 1/f^2^ and broad peaks typically observed in the alpha and beta bands (Fig. 1; Supplementary Fig. 3). There were only small differences observed between WS and N1, and none between WA and S (Fig. 2). By contrast, transitions into states N2 and U were characterized by large band- and region-specific changes in PSDs. As expected, N2 sleep was characterized by a widespread increase in delta power (see Fig. 2a). Of note, increases in alpha power in N2, as might be expected due to sleep spindles (Andrillon et al., 2011), were not consistent across subjects. Loss of responsiveness under anesthesia (U) was associated with large increases in delta power within PFC and sensorimotor areas, and a selective increase in alpha power in PFC (see Fig. 2b), consistent with previous observations (Purdon et al., 2013).

### 3.4 Changes in functional connectivity under sleep and anesthesia

Functional connectivity was assayed using the debiased weighted phase lag index (wPLI) (Vinck et al., 2011). As ECoG power spectra featured peaks in the alpha band, we focused on alpha-band wPLI, but presented analyses of functional connectivity in other canonical frequency bands as well. Like other phase-related measures, wPLI can be sensitive to uncorrelated noise (Vinck et al., 2011), leading to correlations with spectral power. However, in the dataset presented here power did not exhibit an appreciable correlation with wPLI residuals (mean across patients R^2^ = 0.02, maximum R^2^ = 0.04) after accounting for state, indicating that spectral power changes did not contribute substantially to our measure of functional connectivity.

Adjacency matrices were computed first for each pair of recording sites (Fig. 3a), then transformed into ROI-based adjacency matrices (Fig. 3b), from which chord connectivity plots were created (Fig. 3c). Single-subject examples of chord connectivity plots for delta, alpha and gamma bands across arousal states during sleep and anesthesia are shown in Supplementary Figures 4 and 5. Qualitatively, in the wake states (WA, WS) alpha-band connectivity was dominated by connections within and between the temporal and parietal lobes in all five subjects (Fig. 4). This pattern was largely preserved in N1, REM and S states. By contrast, for N2 and U, alpha-band connectivity showed a shift to connectivity within prefrontal ROIs and between prefrontal cortex and select ROIs, including HGPM, insula, gyrus rectus and PMC (see Fig. 4, third column). More modest changes in connectivity were observed in other frequency bands (Supplementary Fig. 6). In the three subjects in whom N3 sleep was observed, the shift in alpha-band connectivity was even more pronounced in N3 compared to N2 (Supplementary Fig. 7).

### 3.5 Common neural signature of functional connectivity changes in sleep and anesthesia

A striking transition boundary in the alpha-band connectivity patterns between two sets of arousal states: [WS, N1, REM, WA, S] and [N2, N3, U] is apparent in the chord connectivity plots. Differences in the degree of conscious experience in these two sets suggest a functional boundary as well: the first set comprises states in which subjects are responsive (WS, WA, S), or have high incidence of reportable conscious experience within the context of dreaming (N1, REM), while the second set comprises states in which subjects are unresponsive and have low incidence of reportable conscious experience (Eer et al., 2009; Leslie et al., 2009; Siclari et al., 2013). To quantify these observations, changes in connectivity with arousal state were measured using the differences between un-thresholded ROI × ROI adjacency matrices. Specifically, the magnitude of the difference in connectivity between states J and K was computed as *d*_J,K_ = ∥**A**_J_ − **A**_K_∥, where **A** is the adjacency matrix for that state and ∥**M**∥ is the operator norm of the matrix **M** (see Methods). Using this metric, functional connectivity was evaluated within each experiment (sleep, anesthesia) to test the hypothesis that differences across the transition boundary (sleep: *d*_N1,N2_ and *d*_REM,N2_; anesthesia: *d*_S,U_) were larger in magnitude than differences that do not cross that boundary (sleep: *d*_WS,N1_, *d*_WS,REM_; anesthesia: *d*_WA,S_). Mean effect sizes across subjects (mean Cliff’s delta, δ◻, see methods) are reported and a permutation test was performed to estimate how chance arrangements of the data compare to the actual differences observed. We found that within the alpha-band, *d*_WS,N1_ was significantly smaller than *d*_N1,N2_ (δ◻ = 0.38, *p* = 0.00013), as was *d*_WA,S_ compared to *d*_S,U_ (δ◻ = 0.70, p = 0.046). Additionally, *d*_WS,REM_ was significantly smaller than *d*_REM,N2_ (δ◻ = 0.25, *p* = 0.0025). Comparable results (i.e. both *d*_WS,N1_ < *d*_N1,N2_ and *d*_WA,S_ < *d*_S,U_ significant) were not found within delta and gamma bands (Supplementary Fig. 6; Supplementary Table 4).

Further support for a transition boundary distinguishing alpha-band connectivity profiles was provided by classification analysis (Fig. 5a). Rather than starting with the average connectivity profiles, as in the difference norms analysis above, the classification analysis was based directly on the minute-by-minute connectivity matrices measured during the overnight sleep experiment. The classifier was trained on data segments from two states appearing to fall on either side of the boundary, WS and N2, and then tested on data segments from all arousal states. We used a logistic weighting function to assign a value between 0 (‘N2-like’) and 1 (‘WS-like’) to each segment. For this analysis, adjacency matrices were calculated from shorter (10-second) segments of data to provide a larger dataset on which to train the classifier, and the analysis was performed on each subject separately. As expected, median prediction scores on N2 and WS were highly skewed toward 0 and 1, respectively (N2: 0.10; WS: 0.90). Separation in median prediction score for N2 and WS segments was greater for alpha (difference of medians = 0.80) compared to other frequency bands (delta, difference of medians = 0.54; gamma, difference of medians = 0.49). N3 data were classified as ‘N2-like’ (median logistic prediction score = 0.12). Importantly, both N1 and REM tended to be classified as ‘WS-like’ (median logistic prediction score = 0.68 and 0.56, respectively). These results were generally consistent across the five subjects (Supplementary Fig. 8).

The similarities between connectivity profiles measured during sleep and anesthesia (i.e. between WS and WA, between N1 and S, and between N2 and U; Fig. 4) suggest a commonality in the mechanisms governing transitions between arousal states in the two experiments. The hypothesis that certain pairs of states in sleep and anesthesia can be considered ‘equivalent’ (i.e. WS and WA, N1 and S, N2 and U) was tested by comparing the distances between alpha-band connectivity profiles measured in equivalent states with those measured in states hypothesized to be ‘non-equivalent’ (i.e. on opposite sides of the transition boundary in Figure 4). Thus, *d*_Equiv_ (i.e. *d*_WS,WA,_ *d*_N1,S_ and *d*_N2,U_) were compared to *d*_Non-equiv_ [i.e. mean (*d*_WS,U_,*d*_WA,N2_), mean (*d*_N1,U_, *d*_S,N2_) and mean (*d*_N1,U_, *d*_S,N2_), respectively]. We found that *d*_WS,WA_ and *d*_N1,S_ were significantly smaller than their corresponding *d*_Non-equiv_ (δ◻ = 0.22, *p* = 0.0022 and δ◻ = 0.23, *p* = 0.00076, respectively) but *d*_N2,U_ was not (δ◻ = 0.14, *p* = 0.31). These data indicate similarity in alpha-band connectivity profiles observed during N1 sleep and sedation. Comparable results (i.e. both *d*_WS,WA_ and *d*_N1,S_ significantly smaller than their corresponding *d*_Non-equiv_) were not found within delta and gamma bands (Supplementary Fig. 6; Supplementary Table 4).

Classification analysis also provided support for the idea that connectivity profiles under sleep and anesthesia overlap. Here, classifiers trained on WS and N2 data from the sleep experiment (see Fig. 5a) were applied to anesthesia data (Fig. 5b) in order to determine whether the transition boundary observed during sleep generalized to changes in arousal state under anesthesia. The classifiers tended to assign WA and S segments to the WS-like category (median logistic prediction score = 0.68 and 0.55, respectively) and assigned U segments with high probability to the N2-like category (median logistic prediction score = 0.11). Taken together, the results of these two analyses suggest substantial overlap in connectivity profiles between ‘equivalent’ sleep and anesthesia arousal states.

### 3.6 Regional distribution of functional connectivity strength across arousal states

The changes in regional distribution of connectivity across the transition boundary, i.e. the shift from temporo-parietal to prefrontal connectivity, were strikingly similar in the sleep and anesthesia experiments (see Fig. 4). Boly and colleagues (Boly et al., 2017) presented evidence that the neural correlates of consciousness correspond primarily to activity in the ‘back’ of the brain, specifically involving broad regions in the temporal, parietal and occipital lobes, and excluding regions in the frontal lobe. Motivated by this perspective, we quantified the differences in regional connectivity observed across arousal states in the current study. We divided ROIs into two groups: a posterior group that included all temporal, parietal and occipital ROIs, and an anterior group that included all frontal ROIs. We then compared the mean alpha-band wPLI across all pairs of recording sites within each group, and calculated a regional bias in connectivity as the difference between within-anterior and within-posterior connectivity. Figure 6a shows the bias in connectivity, with biases toward within-posterior connectivity indicated by negative values and within-anterior by positive values. There was a shift from posterior and towards anterior connectivity with reduced arousal in both sleep [change in regional bias from N2-N1 δ◻ = 0.69, *p* < 0.0001; N2-WS δ◻ = 0.72, *p* < 0.0001] and anesthesia (S-WA δ◻ = 0.98, *p* = 0.0011; U-S δ◻ = 1.0, *p* = 0.00037; U-WA δ◻ = 1.0, *p* < 0.0001). The shift from WS to N1 was not significant (N1-WS δ◻ = 0.34, *p* = 0.093). REM was different from N2 (N2-REM δ◻ = 0.81, *p* < 0.0001) but not significantly different from wake (REM-WS δ◻ = 0.23, *p* = 0.30). Thus, the data indicate that alpha-band connectivity in WS versus N2 and in WA versus U exhibits a similar shift from connectivity within posterior towards connectivity within anterior regions.

Finally, disruption in long-range cortico-cortical connectivity has been noted upon LOC during sleep and anesthesia in several studies (Boly et al., 2012a; Lee et al., 2013b; Ranft et al., 2016; Spoormaker et al., 2010), though these findings have been challenged by other studies (Boly et al., 2012b; Lee et al., 2017; Monti et al., 2013; Murphy et al., 2011). To investigate this issue in the dataset presented here, we assayed the state-dependence of long-range alpha-band connectivity by measuring wPLI across the most distant pairs of recording sites, defined as highest quartile of Euclidean distances in each subject (Fig. 6b). We found no evidence for a decrease in long-range functional connectivity, observing a rather modest increase in N2 and U relative to wake (N2-WS δ◻ = 0.56,, *p* < 0.0001; U-WA δ◻ = 0.74,, *p* = 0.0061) and N1/S (N2-N1 δ◻ = 0.63,, *p* < 0.0001; U-S δ◻ = 0.86,, *p* = 0.0056). We did not find significant changes in long-range connectivity between WS and N1 (N1-WS δ◻ = −0.19,, *p* = 0.15) or WA and S (S-WA δ◻ = −0.05,, *p* = 0.68), but long-range connectivity was reduced in REM (R-WS δ◻ = −0.69,, *p* = 0.00041).

## 4. Discussion

The search for reliable biomarkers of LOC/ROC is of great scientific interest and clinical relevance for anesthesia (Drummond, 2000) as well as for diagnosis and prognosis of disorders of consciousness (Bayne et al., 2017; Bernat, 2017). Here, we leveraged a unique opportunity to obtain intracranial electrophysiological recordings from neurosurgery patients both during natural sleep and under propofol anesthesia. We found that different arousal states were associated with distinct patterns of functional connectivity. This association was similar for sleep and anesthesia, suggesting that cortical network configuration could index changes in consciousness.

### 4.1 ROI- and band-specific effects of sleep and anesthesia on power spectral density

A practical biomarker of conscious vs unconscious state must generalize to multiple settings where LOC is encountered, including sleep and general anesthesia. Previous attempts to use band-specific power to distinguish arousal states under general anesthesia have been largely unsuccessful (Otto, 2008; Struys et al., 1998). This difficulty likely stems from agent-specific changes in power spectra, for example differing between propofol, ketamine and dexmedetomidine anesthesia (Mashour, 2020). The changes that we observed during natural sleep, specifically widespread increases in spectral power in the delta band (see Fig. 2a), are hallmarks of N2 and N3, but not N1, sleep (Prerau et al., 2017; Steriade et al., 1993). In contrast to observations during natural sleep, under propofol anesthesia we observed region-specific (not global) increases in delta power (see Fig. 2b), and increases in frontal alpha power (see Fig. 2b). These observations under propofol are consistent with previous reports (Chennu et al., 2016; Feshchenko et al., 2004; Ni Mhuircheartaigh et al., 2013; Purdon et al., 2013; Supp et al., 2011; Tinker et al., 1977; Wang et al., 2014) and some have suggested that changes in frontal alpha and delta power are reliable indicators of loss of consciousness under propofol (Purdon et al., 2013). However, a recent study using the isolated forearm technique challenges the reliability of such an approach (Gaskell et al., 2017). Consistent with the latter findings, changes in power in the present study did not consistently distinguish N1 from N2 and S from U (see Fig. 2). In addition, these changes across arousal states were not consistently paralleled by changes in connectivity. For example, alpha power did not consistently increase in N2 compared to WS and N1 states, yet this band exhibited the most prominent connectivity changes observed during sleep (see Fig. 2, Fig. 4). Conversely, although the transition to N2 and N3 sleep was characterized by an increase in delta power in multiple ROIs, connectivity within and across these ROIs did not undergo a comparable degree of reorganization (see Fig, 2, Supplementary Fig. 6a). These results indicate that the observed changes in connectivity do not merely follow changes in power and instead reflect functional reorganization of cortical networks. The absence of meaningful correlations between connectivity and power (see Results) further support this idea.

### 4.2 Changes in connectivity during sleep and anesthesia

The sharing of information between cortical regions is a critical element in theories of consciousness and brain function (Dehaene and Changeux, 2011; Friston, 2005; Tononi et al., 2016). Altered cortical connectivity observed during sleep and anesthesia has been interpreted within this theoretical context to explain reduced awareness upon LOC (Alkire et al., 2008; Mashour and Hudetz, 2017). Although there have been studies that examined functional connectivity during sleep and anesthesia (Boly et al., 2012a; Boly et al., 2012b; Lee et al., 2017; Lee et al., 2013b; Murphy et al., 2011; Ranft et al., 2016; Spoormaker et al., 2010), no previous study has directly compared the two in the same subjects. Of particular relevance is the study by Murphy et al. (Murphy et al., 2011) that examined changes in neural activity during sleep and anesthesia. However, that study utilized data from two different sets of subjects and did not compare changes in functional connectivity between the two data sets. A recent study in human volunteers that did measure changes in functional connectivity patterns derived from fMRI during transitions in arousal state found substantial differences between sleep and propofol anesthesia (again, imaged in two different groups of subjects) (Li et al., 2018). Interestingly, the latter study found that cortical changes during NREM sleep were confined to frontal cortex, while changes under propofol anesthesia were widespread. Here, measuring ECoG-derived functional connectivity in the same subjects during sleep and anesthesia, we found substantial overlap in the regional changes in functional connectivity during transitions in arousal state.

We observed consistent and pronounced changes in connectivity upon transitions into N2 and U, specifically increased connectivity within and between anterior (frontal) brain regions, as has been observed using electrophysiological measures previously under propofol anesthesia (Purdon et al., 2013; Supp et al., 2011), and reduced connectivity elsewhere. What is novel about the results presented here is the degree of overlap between changes in connectivity profiles across arousal states in sleep and anesthesia, including a pronounced transition boundary between N1 and N2 and between S and U (Fig. 4). On a superficial level, one might expect some overlap in arousal states, and thus in the changes upon transitions between arousal states, during sleep and anesthesia, yet differences are expected as well. For example, WS and WA are both wake states, but disparities in the time of day of the recordings (overnight versus morning), the behavioral state of the subject (e.g. WA was just prior to major surgery) and environment (monitoring suite versus operating room) could result in substantial differences in cortical network organization. Similarly, although both N2 and U are unresponsive states with low probability of reportable conscious experience, differences in brain state due to the presence of the anesthetic agent versus endogenous sleep factors might result in distinct brain connectivity patterns.

Previous studies of the incidence of dreaming and conscious experience under anesthesia suggest that the observed transition boundary may reflect entry into and out of conscious states. Specifically, on one side of the boundary are states in which subjects are likely having conscious experiences, i.e. responsive (WS, WA, S) or dreaming frequently and vividly with high incidence of reportable conscious experience (REM, N1). On the other side are arousal states in which subjects are unlikely to be having conscious experiences, i.e. unresponsive and with low incidence of reportable conscious experience (Leslie et al., 2009; Siclari et al., 2013). This boundary was observed both with difference norms and classification analyses applied to the ROI-by-ROI adjacency matrices (see Fig. 4, 5) and with the analysis of intra-regional and long-range connectivity (see Fig. 6). However, even though connectivity patterns during propofol sedation (S) generally aligned with other conscious states, both the classification and intra-regional connectivity analyses were consistent with fluctuations in arousal level in this state (see Fig. 5b, 6a).

A recent essay on the neural correlates of consciousness (NCC) suggests an interesting interpretation of these changes in connectivity. Boly and colleagues (Boly et al., 2017) presented evidence from lesion studies and from experiments utilizing serial awakening during sleep to argue that the “full NCC”, that is the collection of all regions underlying specific contents of consciousness, comprises large portions of the parietal, occipital, and temporal lobes, whereas frontal lobe structures underlie functions associated with, but not necessary for, those conscious contents. The regions within the full NCC are most closely associated with sensory awareness, and thus would underlie the internal generative models central to theories of predictive processing and the mismatch detection and message passing functions critical to those schemes (Friston, 2005). Alpha-band power and phase synchronization in particular are associated with feedback connectivity in the visual cortical hierarchy (van Kerkoerle et al., 2014). Thus, it is possible that the shift in cortical connectivity from predominantly temporo-parieto-occipital (posterior) to frontal (anterior) upon LOC may reflect a reduction in predictive processing during states of reduced consciousness. This is consistent with the finding that anterior alpha synchronization of EEG in response to propofol correlates with disrupted sensory processing in human volunteers (Supp et al., 2011).

Although clinical considerations precluded electrode coverage of the thalamus, previous studies suggest that some of the changes in cortico-cortical connectivity observed in this study could be driven by altered thalamo-cortical synchronization (Saalmann et al., 2012). For example, the increased thalamo-cortical synchronization observed during sleep spindles (Andrillon et al., 2011) and during propofol anesthesia (Flores et al., 2017) may have a similar effect on functional connectivity within frontal cortex, as suggested by computational studies (Vijayan et al., 2013). However, the observations that the frontal shift in alpha-band connectivity was even more pronounced in N3 than it is in N2 (Supplementary Fig. 7), even though spindles are less common in N3 (Andrillon et al., 2011), and that significant changes in alpha power were not observed during sleep (see Fig. 2a), suggest that the changes in alpha-band connectivity were unlikely driven solely by sleep spindle activity.

The disintegration of cortical networks observed upon LOC during sleep, anesthesia and coma (Alkire et al., 2008) has been ascribed to disrupted long-range connectivity. For example, several reports suggest reduced resting-state cortico-cortical (fronto-parietal) feedback connectivity under a variety of anesthetic agents, including propofol (fMRI: Boly et al., 2012a; Ranft et al., 2016; EEG: Lee et al., 2013b), consistent with results using invasive electrophysiological recordings in rodent models (Imas et al., 2005; Raz et al., 2014). Disrupted long-range resting-state functional connectivity has also been reported in fMRI studies during NREM sleep (Spoormaker et al., 2010) and anesthesia (Ranft et al., 2016). However, other studies have shown no differences in changes in short-versus long-range connectivity (fMRI: Monti et al., 2013), or even increases in long-range connectivity during anesthesia (fMRI: Murphy et al., 2011; EEG: Lee et al., 2017) and sleep (fMRI: Boly et al., 2012b). Similarly, in the present study, we saw little evidence for decreases specifically in long-range connections (see Fig. 6b). The reasons for the diverse findings of the effects on connectivity are unclear. It is possible that the dynamics and heterogeneity of the resting state cortical network contribute to this diversity. For example, network configuration prior to LOC has been shown to influence observed changes in connectivity during sleep (Wilson et al., 2019). Application of methods to these data that can characterize connectivity at finer temporal resolution may address this issue.

### 4.3 Caveats and limitations

The key limitations of this study are the small number of participants (*n* = 5), and that the subjects had a neurologic disorder, and thus may not be entirely representative of a healthy population. These caveats are inherent to all human intracranial electrophysiology studies. Our statistical methods focused on within-subject comparisons between states and should be generalized with caution. However, results were consistent across subjects who all had different clinical histories of their seizure disorder, antiepileptic medication regimens, and seizure foci. Recordings from cortical sites confirmed to be seizure foci were excluded from analyses. Finally, all subjects participated in multiple additional research protocols over the course of their hospitalization, including a range of behavioral tasks. Behavioral and neural data obtained in these other experiments were examined for consistency with a corpus of published human intracranial electrophysiology data (reviewed in Nourski, 2017). None of the subjects exhibited aberrant responses that could be interpreted as grounds for caution in inclusion in this study.

The motivation for exploring changes in connectivity across arousal states is to elucidate the neural underpinnings that define these states. We note, however, that the arousal states as defined in this study are likely non-uniform regarding consciousness. For example, healthy adults are able to report on conscious experience (i.e. dreaming) about 40% and 20% of the time in N2 and N3 sleep (Siclari et al., 2013). Dreaming also occurs under propofol anesthesia in about 20% of patients (Leslie et al., 2009). This suggests that differences in brain connectivity between the conscious and unconscious states may be even greater than those reported here, had it been possible to reliably distinguish dreaming vs. non-dreaming states in our data set.

We also note the challenges in assessing awareness under anesthesia, and specifically the delicate balance between interrogating a subject’s awareness and changing the state of their arousal with that interrogation. The approach employed here, the OAA/S, is considered the gold standard for assessing awareness in the perioperative setting (Chernik et al., 1990), and it has been cross-validated using EEG-based measures such as BIS (Vanluchene et al., 2004). The BIS values recorded in the current study corresponded well to those associated with wake, sedated and unconscious states in previous reports (Vanluchene et al., 2004). Importantly, we did not observe consistent increases in BIS values post-OAA/S assessments compared to pre-OAA/S assessments (see Supplementary Fig. 2), indicating that our assessments likely did not alter the arousal state of the subjects.

### 4.4 Functional significance and future directions

The results presented here have broad implications for understanding the neural mechanisms associated with loss of consciousness and for better understanding and differential diagnosis of disorders of consciousness. We demonstrate a transition boundary in profiles of functional connectivity that separates states of different levels of consciousness. Phase synchronization is postulated to mediate rapid communication of conscious content over multiple spatial scales in cortex, contributing importantly to the rich repertoire of human behavior that characterizes conscious states (Fries, 2015). The finding that changes in functional connectivity based on phase synchronization indexes arousal state similarly in both sleep and anesthesia motivates further exploration of the changes in brain activity and connectivity common to changes in consciousness. These findings have practical clinical ramifications as well. Connectivity can be measured non-invasively using EEG or fMRI in patients with disorders of consciousness. Algorithms that track region-specific functional connectivity may provide a basis for noninvasive monitoring of arousal state in patients otherwise inaccessible to standard assessments of arousal based on response to command. Future experiments aimed at exploring in more detail the differences between LOC in sleep and anesthesia, and generalizing to other anesthetic agents such as dexmedetomidine and volatile anesthetics, will elucidate further fundamental questions about the nature of consciousness and arousal that remain unresolved.

## Acknowledgements

This work was supported by the National Institutes of Health (grant numbers R01-DC04290, R01-GM109086, UL1-RR024979). We are grateful to Jess Banks, Haiming Chen, Phillip Gander, Christine Glenn, Bradley Hindman, Matthew Howard, Rashmi Mueller, Ariane Rhone, Robert Sanders, Beau Snoad, Mitchell Steinschneider, Deanne Tadlock, and Thoru Yamada for help with data collection, analysis, and comments on the manuscript.

## Author Contributions

M.I.B and K.V.N. designed the experiments. M.I.B., C.K.K., H.K. and K.V.N. collected the data. M.I.B., B.M.K., C.M.E., D.I.C., C.K.K., M.E.D. and K.V.N. analyzed the data. M.I.B., B.M.K. and K.V.N. drafted and revised the manuscript.

