## Supplementary Online Material for "Cortical functional connectivity indexes arousal state during sleep and anesthesia"

***Corresponding author:**

Matthew I. Banks, Ph.D.

Professor

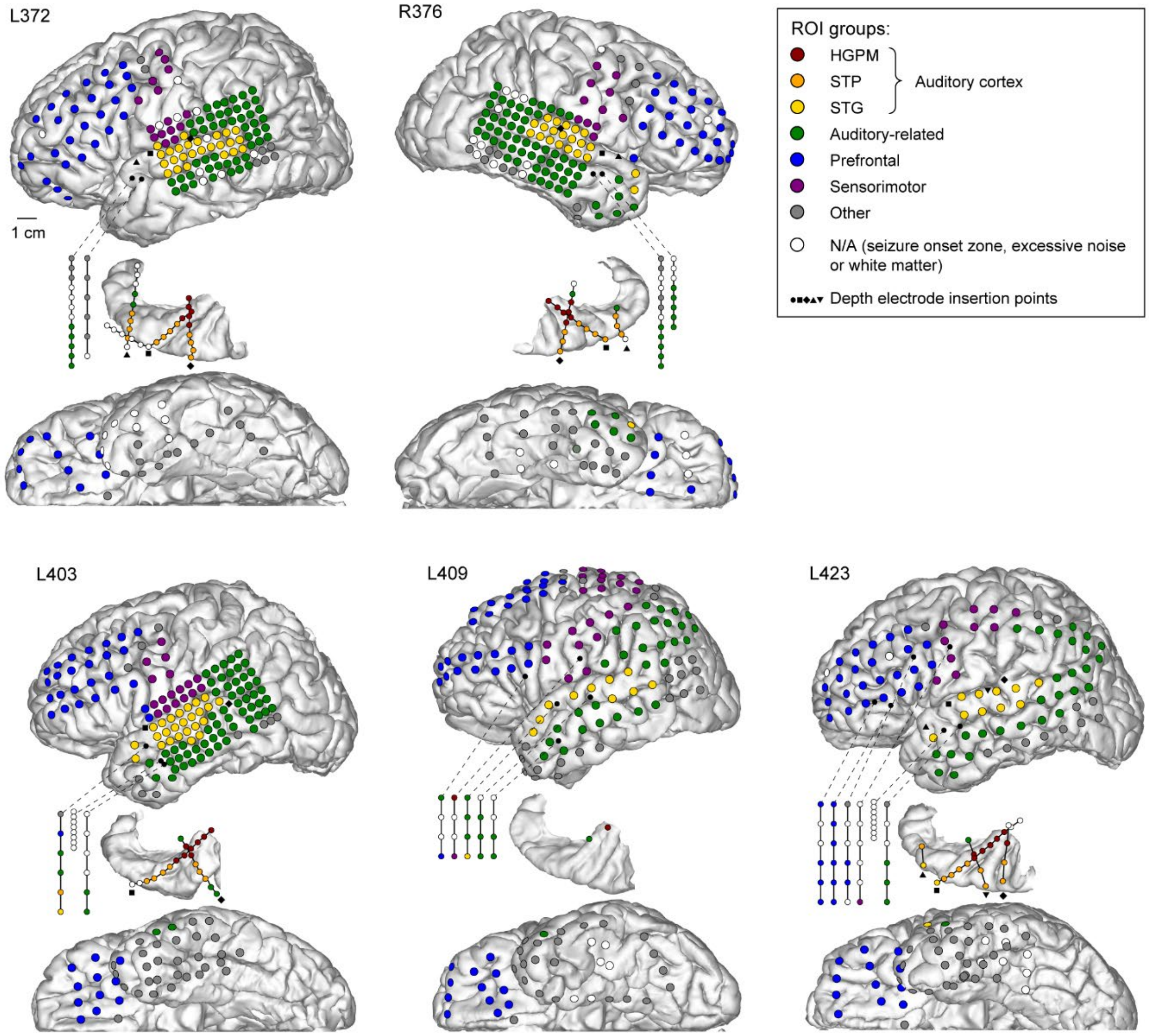


**Supplementary Fig. 1: Electrode coverage in all subjects.** For each subject, the three views are (top-to-bottom): lateral view of cerebral hemisphere, top-down view of superior temporal plane and bottom-up view of cerebral hemisphere. Depth electrode coverage is shown only for the superior temporal plane; all subjects had additional depth electrode arrays, shown schematically (most medial contacts are on top). Black symbols indicate depth electrode insertion points on the hemisphere lateral surface. In subject L409, the two contacts in the STP (one in insular cortex, the other in HGPM, green and dark red, respectively) were on two of the five depth electrode arrays shown schematically in the panel. See Supplementary Table 2 for total numbers and regional distribution of recording sites.

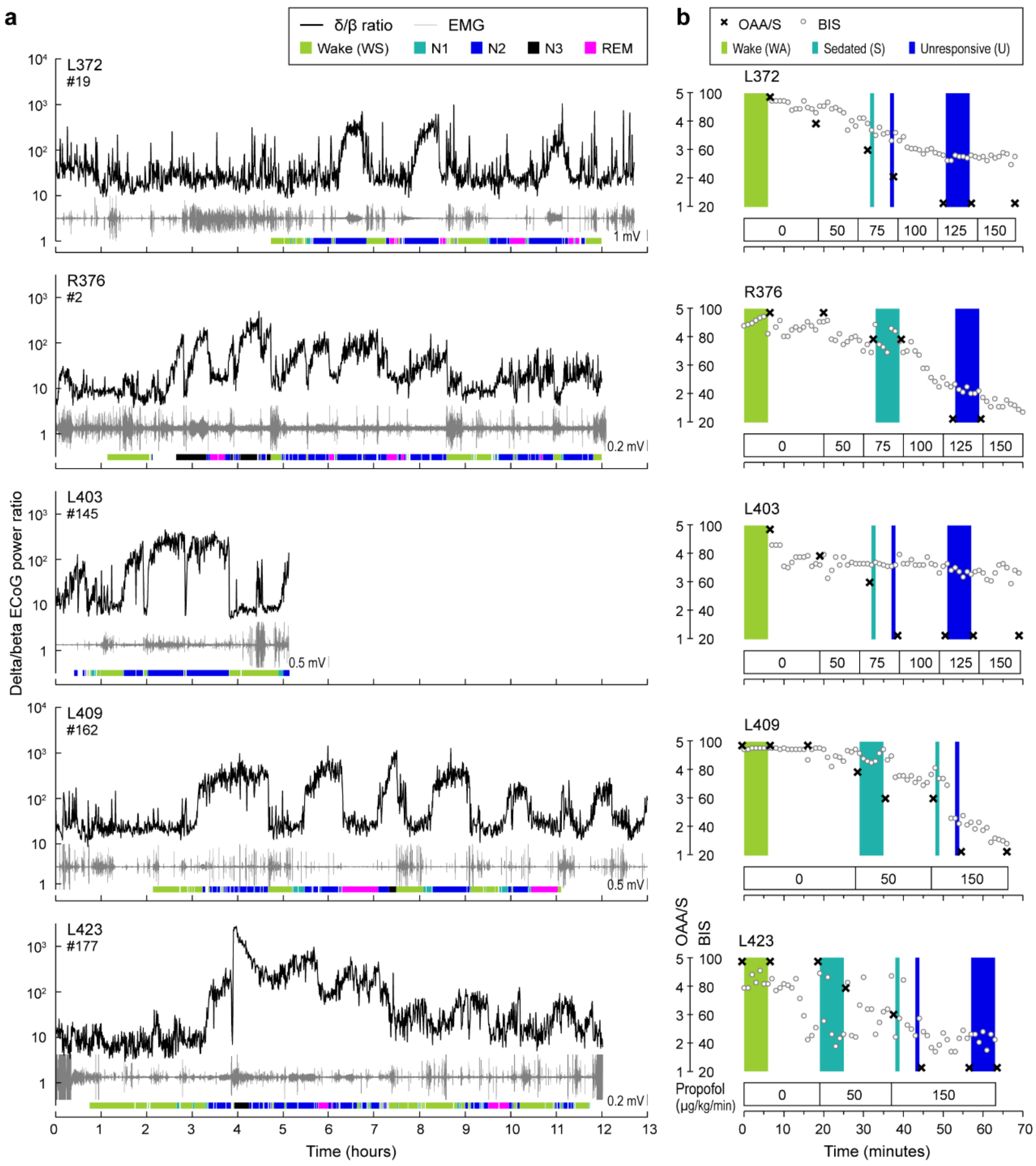


**Supplementary Fig. 2: Experiment time course in all subjects. a,** Overnight sleep time course. For each of the five subjects (rows), delta/beta power ratio for a representative intracranial recording site in TFG, and electromyogram, (EMG) are plotted as functions of time in black and gray, respectively. Color bars underneath highlight wake and different sleep stages, as determined by sleep stage scoring. **b,** Induction of general anesthesia. Observer's Assessment of Alertness/Sedation scores (crosses) and bispectral index (BIS) values (open circles) are plotted as functions of time. Propofol infusion rates are shown underneath each plot. Colored rectangles denote recording epochs corresponding to the three brain states (green: wake; light blue: sedated; dark blue: unresponsive).

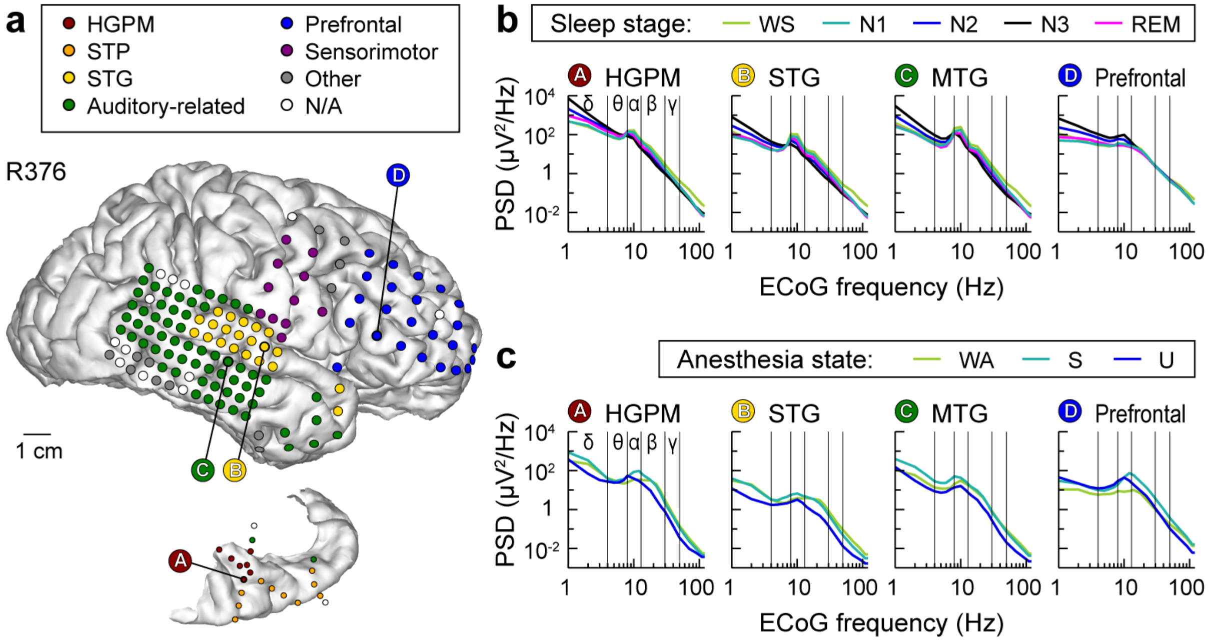


**Supplementary Fig. 3: Electrode coverage (a) and ECoG power spectra during sleep (b) and anesthesia (c).** Exemplary data from subject R376. See caption of Fig. 1 for details.

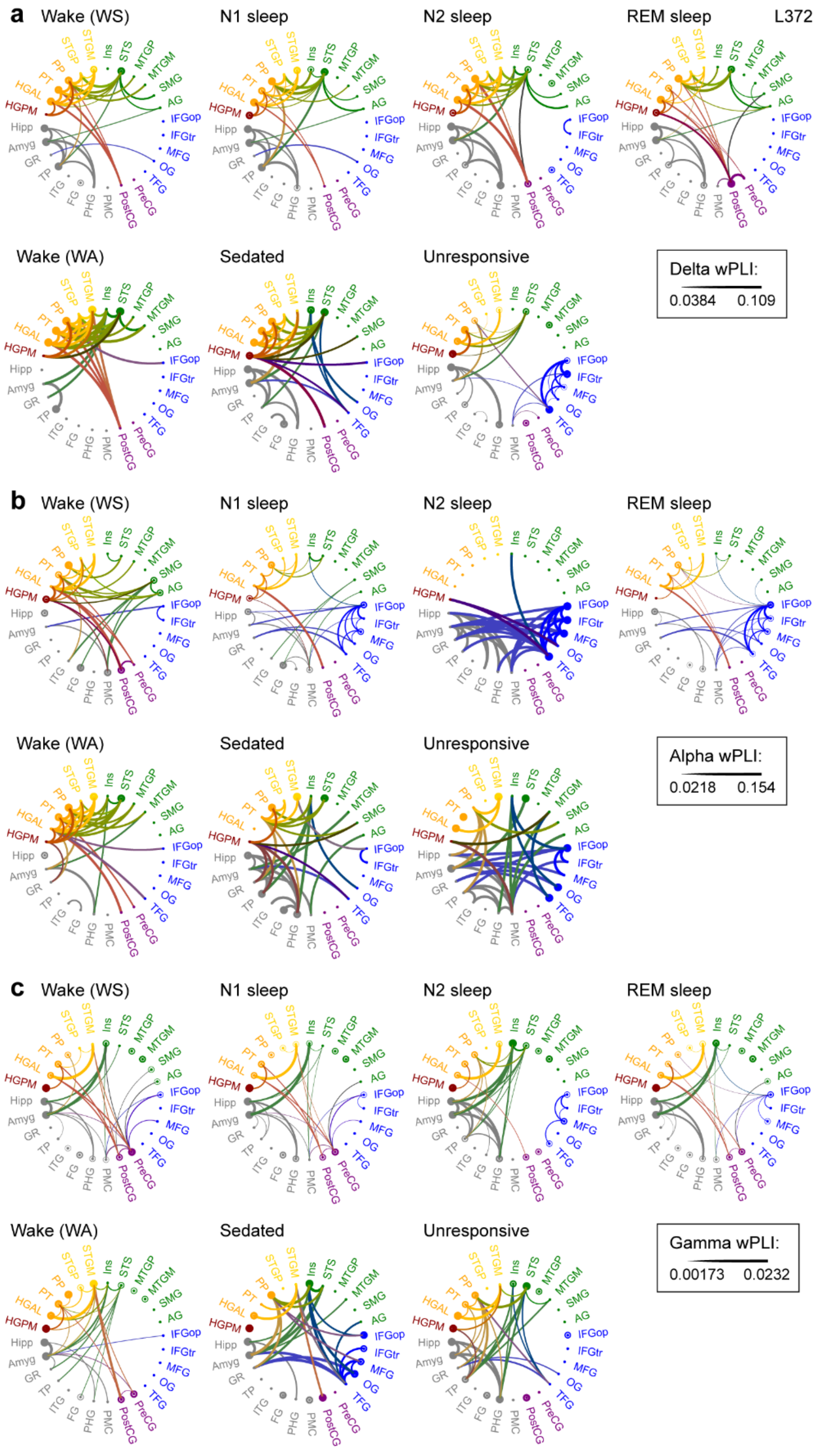


**Supplementary Fig. 4: ROI-based analysis of functional connectivity across brain states in delta (a), alpha (b) and gamma (c) bands.** Exemplary data from subject L372. See caption of **Figure 3c** for details.

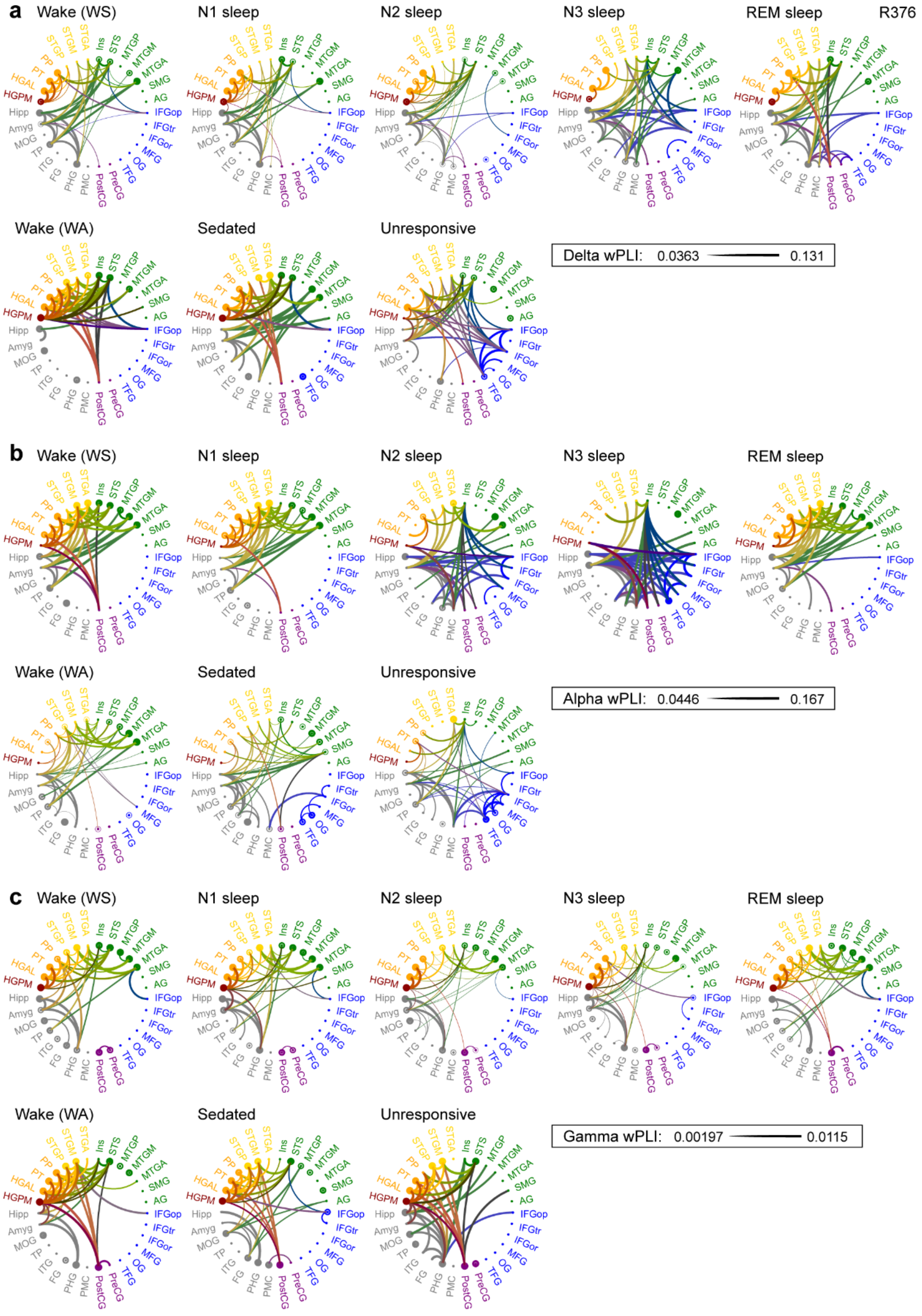


**Supplementary Fig. 5: ROI-based analysis of functional connectivity across brain states in delta (a), alpha (b) and gamma (c) bands.** Exemplary data from subject R376. See caption of **Figure 3c** for details.

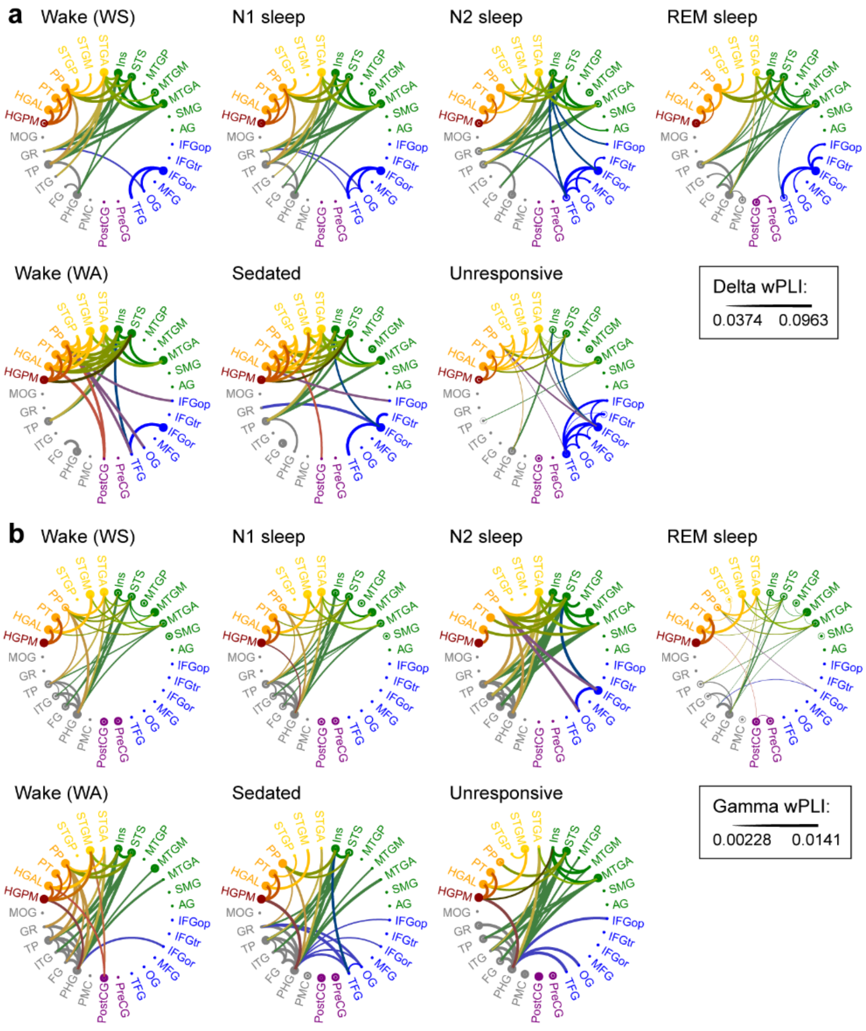


**Supplementary Fig. 6: ROI-based analysis of functional connectivity across brain states in delta (a) and gamma (b) bands.** Data from five subjects. See caption of **Figure 3c** for details.

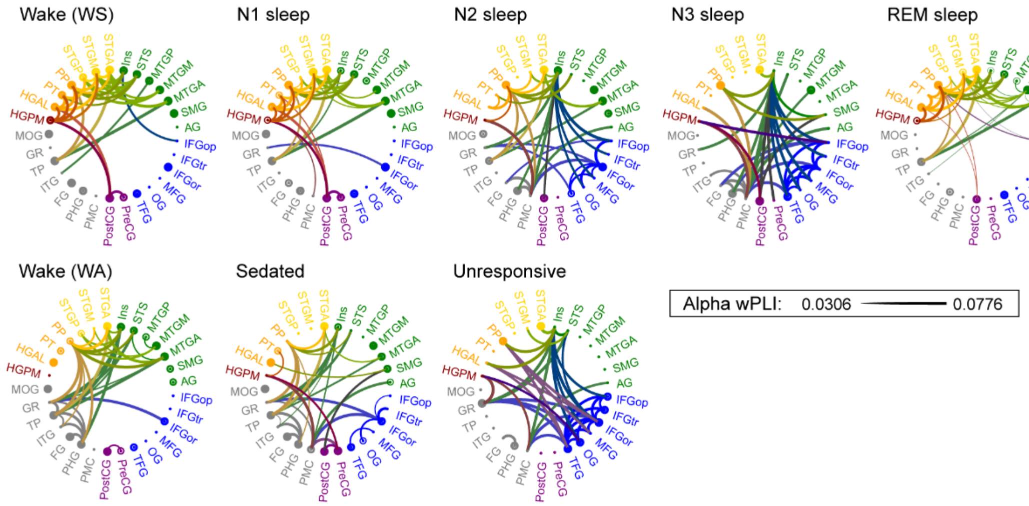


**Supplementary Fig. 7: ROI-based analysis of functional connectivity across brain states in alpha band**.

Data from three subjects with N3 sleep data. See caption of **Figure 3c** for details.

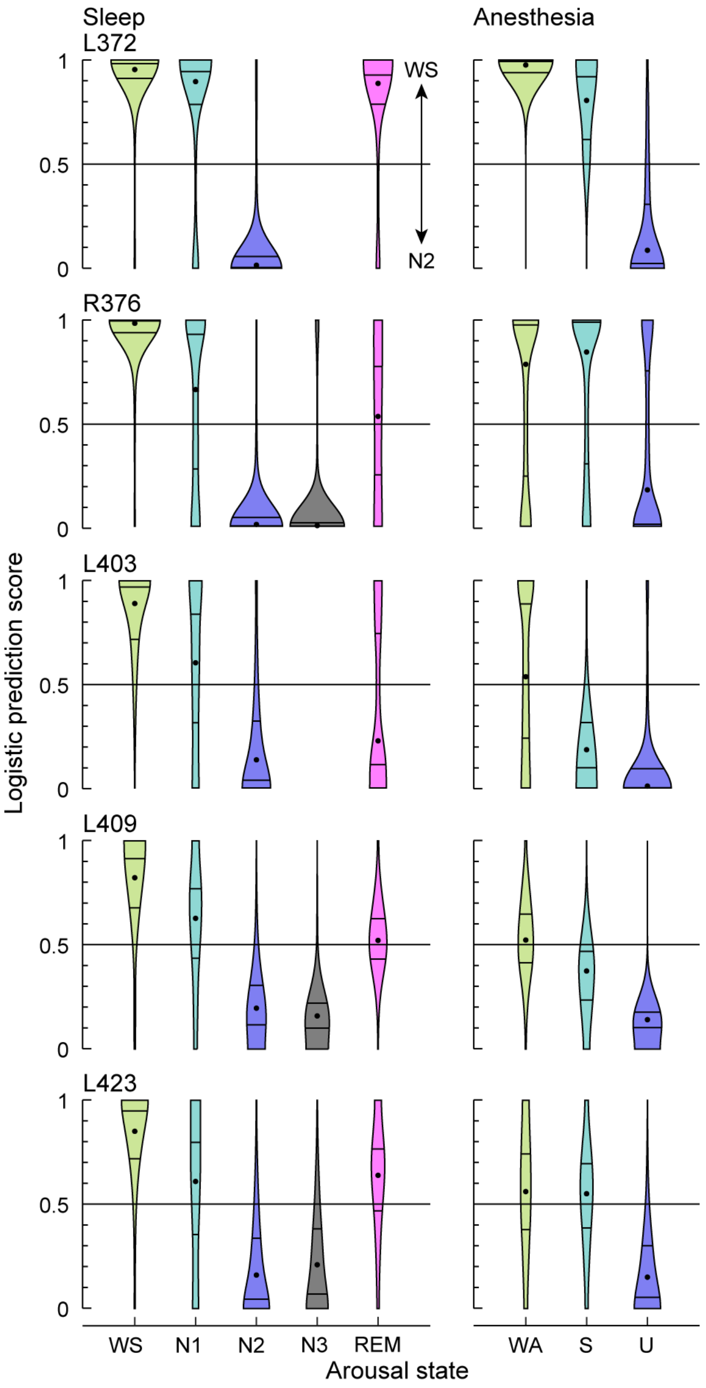


**Supplementary Fig. 8: Classification of data segments in individual subjects.** Logistic prediction distributions for adjacency matrices from sleep and anesthesia arousal states (left and right column, respectively) analyzed by a linear classifier trained on a subset of WS and N2 data. All distributions are scaled to the same total area, and all plots are on the same scale.

**Supplementary Table 1.** Subject demographics.

| Subject[1] | Age | Sex[2] | Seizure focus |
|---|---|---|---|
| **L372** | 34 | M | L temporal pole |
| **R376** | 48 | F | R medial temporal lobe |
| **L403** | 56 | F | L medial temporal lobe |
| **L409** | 31 | F | L medial temporal lobe |
| **L423** | 51 | M | L medial temporal lobe |

[1]Letter prefix of the subject code denotes the side of electrode implantation over auditory cortex, and the side of seizure focus (L = left; R = right). Most subjects had, to varying degrees, bilateral coverage of other regions of the brain.

[2]F = female; M = male

**Supplementary Table 2.** Electrode coverage.

| ROI group | ROI | Number of recording sites | | | | | |
|---|---|---|---|---|---|---|---|
| | | L372 | R376 | L403 | L409 | L423 | Total |
| Auditory core | HGPM | 6 | 7 | 8 | 1 | 7 | 29 |
| STP | HGAL | 4 | 4 | 4 | - | 4 | 16 |
| | PT | 4 | 3 | 3 | - | 4 | 14 |
| | PP | 3 | 3 | 1 | - | 1 | 8 |
| STG | STGP | 23 | 16 | 14 | 6 | 3 | 62 |
| | STGM | 2 | 2 | 9 | 1 | 7 | 21 |
| | STGA | - | 2 | 3 | 3 | 2 | 10 |
| Auditory-related | Ins | 2 | 4 | 3 | 2 | 2 | 13 |
| | STS | 5 | 9 | - | 4 | 3 | 21 |
| | MTGP | 12 | 22 | 28 | 3 | 5 | 70 |
| | MTGM | 6 | 16 | 8 | 4 | 3 | 37 |
| | MTGA | 0 | 6 | 8 | 5 | 1 | 20 |
| | SMG | 21 | 11 | 17 | 6 | 2 | 57 |
| | AG | 9 | 5 | 1 | 12 | 13 | 40 |
| Prefrontal | IFGop | 5 | 2 | 2 | 3 | 5 | 17 |
| | IFGtr | 5 | 3 | 2 | 4 | 7 | 21 |
| | IFGor | - | 1 | 1 | - | 3 | 5 |
| | MFG | 12 | 12 | 16 | 6 | 11 | 57 |
| | SFG* | - | - | - | 18 | 1 | 19 |
| | OG | 7 | 6 | 14 | 12 | 18 | 57 |
| | TFG | 3 | 5 | 1 | 4 | 2 | 15 |
| | CGA* | - | - | - | - | 1 | 1 |
| Sensorimotor | PreCG | 8 | 5 | 11 | 9 | 7 | 40 |
| | PostCG | 6 | 6 | 6 | 11 | 3 | 32 |
| Other | PMC | 2 | 4 | 4 | 3 | - | 13 |
| | PHG | 5 | 2 | 3 | 1 | 4 | 15 |
| | FG | 4 | 2 | 5 | 4 | 2 | 17 |
| | ITG | 6 | 13 | 13 | 7 | 18 | 57 |
| | TP | 3 | 7 | 11 | 11 | 8 | 40 |
| | GR | 1 | - | 2 | 2 | 1 | 6 |
| | SPL* | - | - | - | 2 | 2 | 4 |
| | MOG | - | 3 | - | 7 | 2 | 12 |
| | IOG* | - | - | - | - | 1 | 1 |
| | LG* | - | - | - | 2 | - | 2 |
| | CGM* | - | - | - | 2 | 1 | 3 |
| | Amyg* | 2 | 2 | - | - | - | 4 |
| | Hipp* | 6 | 2 | - | - | - | 8 |
| **Total** | | 172 | 185 | 198 | 155 | 154 | 864 |

*Limited coverage (present in 2 or fewer subjects out of 5)

**Supplementary Table 3.** Summary of sleep data.

| **Sleep stage** | **Recording duration (min.)** | | | | |
|---|---|---|---|---|---|
| | **L372** | **R376** | **L403** | **L409** | **L423** |
| Wake (WS) | 152.5 | 166.1 | 119.4 | 177.0 | 324.1 |
| N1 | 27.5 | 25.0 | 38.5 | 63.2 | 51.9 |
| N2 | 203.1 | 299.8 | 147.9 | 201.7 | 210.7 |
| N3 | - | 79.8 | - | 9.5 | 20.0 |
| REM | 51.6 | 49.5 | 1.0 | 83.0 | 42.9 |
| **Total scored data** | 434.7 | 620.2 | 306.8 | 534.4 | 649.6 |

**Supplementary Table 4.** Norm contrasts for delta and gamma bands.

| Band | Comparison | Cliff's delta (d) | p-value |
|---|---|---|---|
| delta | $d_{WS,N1} < d_{N1,N2}$ | 0.248 | 0.013 |
| | $d_{WA,S} < d_{S,U}$ | 0.230 | 0.212 |
| | $d_{WS,REM} < d_{REM,N2}$ | -0.084 | 0.755 |
| | $d_{WS,WA} < d_{nonequiv}$ | 0.169 | 0.016 |
| | $d_{N1,S} < d_{nonequiv}$ | 0.090 | 0.209 |
| | $d_{N2,U} < d_{nonequiv}$ | 0.001 | 0.883 |
| gamma | $d_{WS,N1} < d_{N1,N2}$ | 0.067 | 0.432 |
| | $d_{WA,S} < d_{S,U}$ | 0.243 | 0.172 |
| | $d_{WS,REM} < d_{REM,N2}$ | 0.016 | 0.457 |
| | $d_{WS,WA} < d_{nonequiv}$ | 0.068 | 0.004 |
| | $d_{N1,S} < d_{nonequiv}$ | -0.100 | 0.034 |
| | $d_{N2,U} < d_{nonequiv}$ | -0.286 | 0.998 |
